# Lipid-mediated regulation of pneumolysin balances vascular injury and bacterial containment during pneumococcal infection

**DOI:** 10.64898/2026.06.23.733954

**Authors:** Martina Kiefmann, Sven Hammerschmidt, Alexander Dietrich, Rainer Kiefmann

## Abstract

Severe infection with *Streptococcus (S.) pneumoniae* is a leading cause of acute lung injury, multiorgan failure, and death despite appropriate antibiotic therapy. A major contributor to tissue damage is pneumolysin (PLY), a cholesterol-dependent pore-forming toxin that is released in large amounts during bacterial lysis. Because antibiotic treatment enhances PLY release, understanding the mechanisms that control PLY-induced host injury is critical for the development of strategies that limit tissue damage without impairing antibacterial defense.

Using isolated perfused lungs and *in situ* two-photon imaging, we show that *S. pneumoniae* induces platelet and leukocyte recruitment, pulmonary edema, perfusion failure, and intravascular bacterial trapping within pulmonary arterioles. These responses required PLY and were associated with endothelial Ca²⁺ influx and plasma membrane permeabilization.

The oxysterol 25-hydroxycholesterol (25-HC) prevented PLY-dependent Ca²⁺ entry, vascular leakage, and perfusion failure by reducing accessible membrane cholesterol through activation of acyl-CoA:cholesterol acyltransferase (ACAT). Unexpectedly, pneumococcal infection suppressed expression of cholesterol-25-hydroxylase (CH25H), the enzyme responsible for 25-HC synthesis, although induction of this pathway would be predicted to protect against toxin-mediated injury. We show that this downregulation results from inhibition of platelet-activating factor (PAF) receptor (PAFR) signaling by pneumococcal phosphocholine-containing cell wall components.

Remarkably, both PAF itself and the anti-phosphocholine antibody TEPC directly inhibited PLY pore formation independently of PAFR, while preserving intravascular coagulation and bacterial trapping. Thus, although 25-HC protects against toxin-mediated vascular injury, its suppression may allow coagulation-dependent bacterial containment, with PAF acting as a compensatory antagonist of PLY.

These findings identify a lipid-based host defense system in which 25-HC and PAF differentially regulate PLY activity to balance tissue protection with bacterial clearance and suggest new therapeutic approaches to limit the harmful effects of antibiotic-induced PLY release during pneumococcal infection.

## Introduction

*Streptococcus (S.) pneumoniae* is a leading cause of community-acquired pneumonia and sepsis and remains associated with high mortality due to acute lung injury and multiorgan failure. Clinical deterioration frequently continues despite appropriate antibiotic therapy, indicating that host tissue damage, rather than bacterial burden alone, is a major determinant of outcome. A major contributor to injury during pneumococcal infection is pneumolysin (PLY), a cholesterol-dependent pore-forming toxin that is released in large amounts during bacterial lysis. Because antibiotic treatment promotes PLY release, understanding how the host controls PLY-induced injury is essential for developing strategies that limit tissue damage without compromising antibacterial defense^1^.

PLY binds to cholesterol-rich plasma membranes (PM), forms transmembrane pores, and induces sustained Ca²⁺ influx in host cells ^2^. In the pulmonary microvasculature, PLY has been linked to endothelial barrier disruption, platelet activation, leukocyte recruitment, and edema formation, resulting in perfusion failure and impaired gas exchange ^3^. Furthermore, intravascular coagulation during pneumococcal infection may contribute to host defense by trapping bacteria within the lung circulation ^4,5^. These observations suggest that the host must balance protection from toxin-induced injury with preservation of intravascular mechanisms required for bacterial containment, but the mechanisms that regulate this balance are poorly understood.

PM lipid composition critically determines susceptibility to cholesterol-dependent cytolysins (CDCs). The oxysterol 25-hydroxycholesterol (25-HC), generated by the enzyme cholesterol-25-hydroxylase (CH25H), reduces the availability of accessible PM cholesterol through activation of acyl-CoA:cholesterol acyltransferase (ACAT), thereby inhibiting PM pore formation by several CDCs ^6–9^. In addition to its antiviral and immunomodulatory functions, 25-HC has been implicated in the regulation of endothelial permeability, inflammation, and coagulation, suggesting that oxysterol-mediated lipid remodeling may represent a host defense mechanism against CDC-induced injury ^10,11^.

Based on these observations, induction of CH25H during pneumococcal infection would be expected to protect the host from PLY-mediated damage. However, whether the CH25H–25-HC pathway is activated or suppressed during pneumococcal infection is unknown, and the functional consequences of its regulation for vascular injury and bacterial clearance have not been determined.

Another pathway that may influence this balance involves the platelet-activating factor (PAF) receptor (PAFR). *S. pneumoniae* express phosphocholine (ChoP)-containing cell wall components that bind to PAFR and interfere with receptor signaling ^12^. Because PAFR activation is linked to intracellular Ca²⁺ regulation, inflammatory activation, and platelet responses ^13^, inhibition of this pathway could affect both lipid metabolism and toxin susceptibility during infection. Whether pneumococcal interaction with PAFR regulates CH25H expression or modifies the host response to PLY has not been investigated.

In the present study, we examined how pneumococcal infection induces pulmonary microvascular injury and how host lipid mediators regulate this response. We show that PLY is required for pneumococcus-induced perfusion failure, leukocyte and platelet recruitment, and edema formation in the lung, and that these responses depend on PLY-induced PM pore formation and Ca²⁺ entry. We further demonstrate that 25-HC protects against PLY-mediated injury by reducing accessible PM cholesterol, but unexpectedly find that pneumococcal infection suppresses CH25H expression through inhibition of PAFR signaling. Finally, we identify PAF itself as a direct antagonist of PLY PM pore formation that preserves intravascular coagulation while limiting vascular injury. These findings reveal a lipid-based host defense mechanism that balances tissue protection with bacterial containment and suggest new strategies to counteract the harmful effects of antibiotic-induced PLY release during pneumococcal infection.

## Results

### Pneumococcal lung injury requires pneumolysin, extracellular Ca²⁺, and is prevented by 25-hydroxycholesterol

Lung injury is characterized by rapid deterioration of pulmonary function associated with microvascular obstruction, edema formation, and inflammatory cell recruitment ^5,14,15^. To investigate the mechanisms linking pneumococcal infection to pulmonary vascular dysfunction, we analyzed microvascular responses to *S. pneumoniae* in isolated perfused mouse lungs using *in situ* two-photon microscopy.

Perfusion of lungs with wild-type *S. pneumoniae* D39 caused marked recruitment of platelets and leukocytes to pulmonary arterioles and capillaries, accompanied by progressive impairment of microvascular perfusion. Within minutes after bacterial exposure, platelet adhesion and leukocyte accumulation increased, followed by a reduced red blood cell velocity, heterogeneous perfusion, intravascular RBC clot formation, and the appearance of non-perfused vascular segments (Fig. 1A–E). In parallel, D39 exposure elevated vascular permeability associated with formation of interstitial and alveolar edema and increased lung wet-to-dry weight ratio and pulmonary artery pressure (Fig. 1F–H).

**Figure 1.**
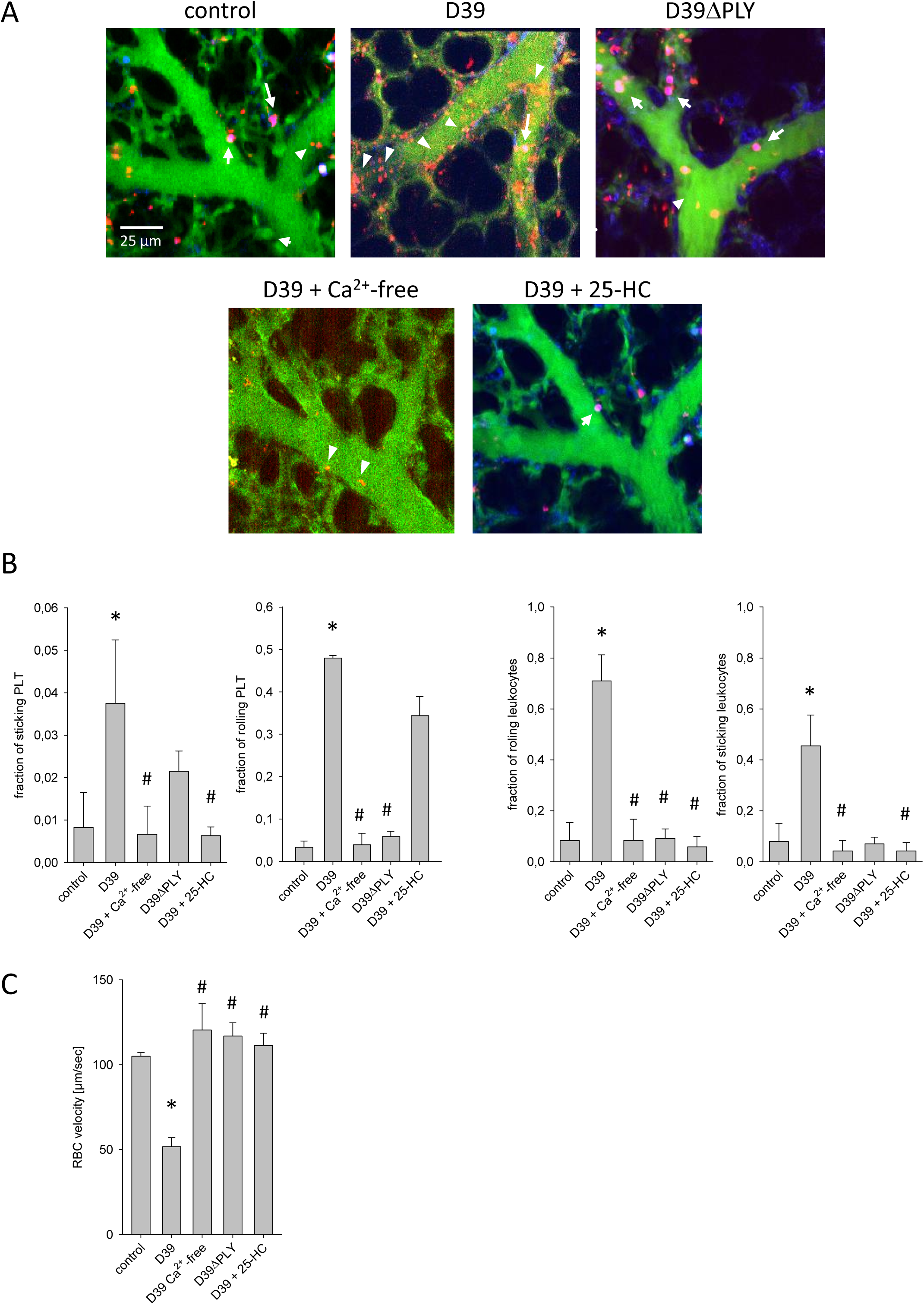
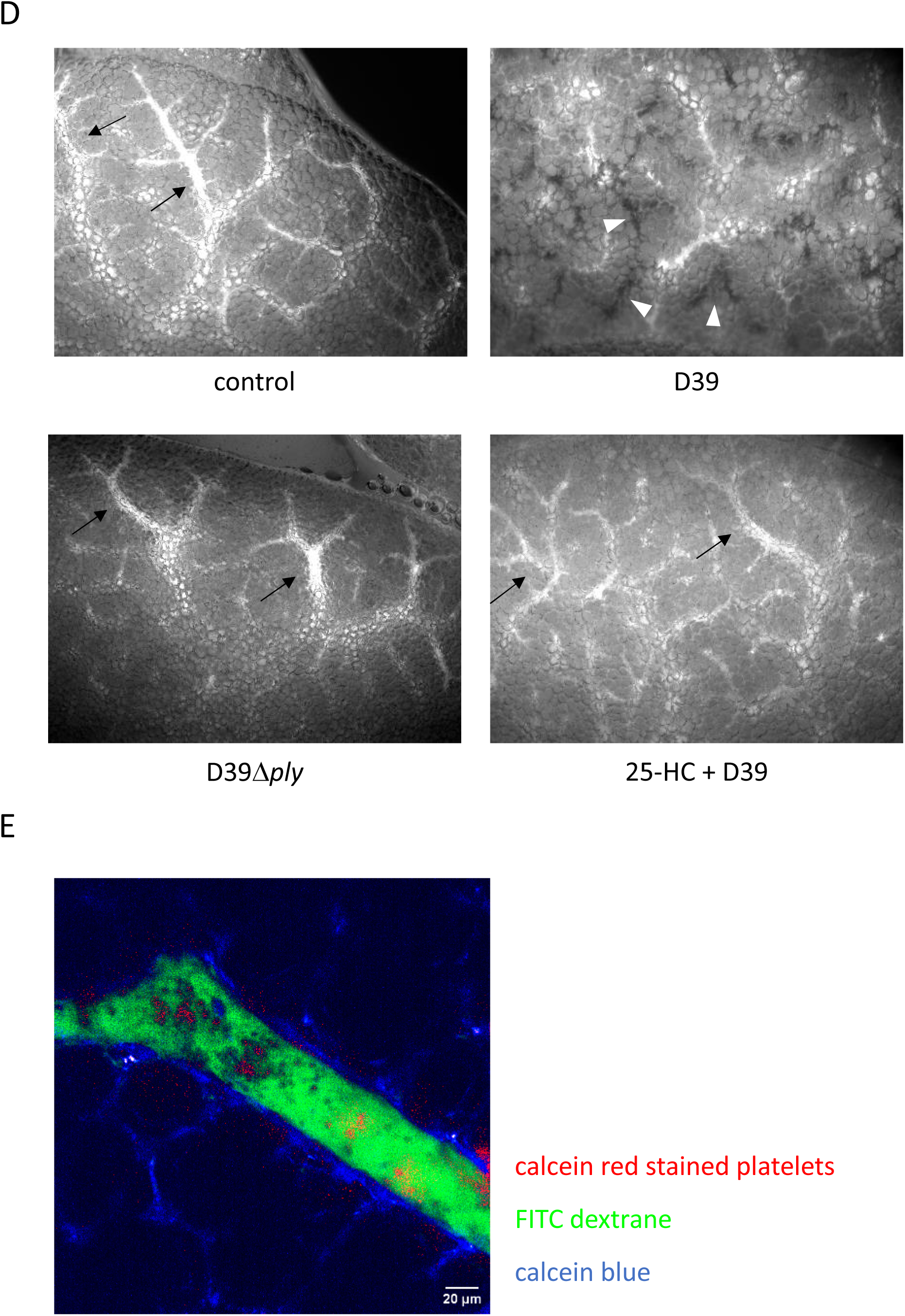
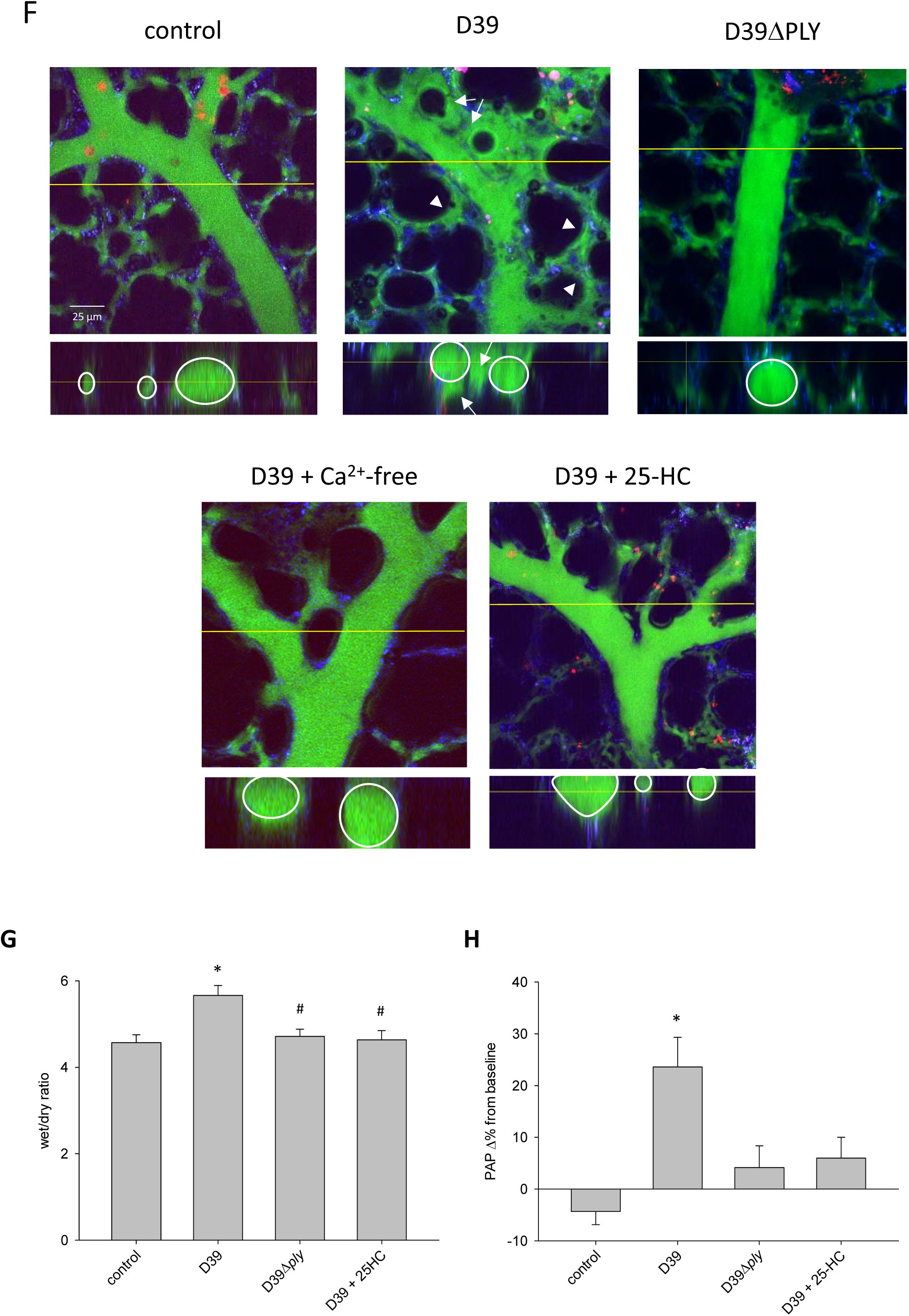
Leukocyte and platelet–endothelial interaction *in situ*. **(A)** Top row: Two-photon microscopy images of subpleural pulmonary arterioles from isolated, buffer-perfused mouse lungs under control condition and after infusion of D39 or D39ΔPLY (5×10⁵ CFU/mL each), in the presence or absence of 25-HC (50 µM) and under Ca²⁺-containing or Ca²⁺-free conditions (as indicated). Vessels were co-perfused with FITC-dextran (green) and calcein red-labeled leukocytes and platelets. Still frames show adherent leukocyte (red, arrows) and platelets (red, arrow heads). **(B)** Quantification of leukocyte– and platelet–endothelial interactions. Fraction of rolling or sticking platelets or leukocytes, as indicated (rolling defined as platelets or leukocytes with reduced velocity or pauses <10 s, sticking defined defined as immobile for >10 s). Data are shown as mean ± SEM. ^#^p < 0.05 vs. D39; *p < 0.05 vs. D39 + 25-HC; n = 3–11. **(C)** Quantification of red blood cell (RBC) velocity under the indicated conditions. Data are shown as Mean ± SEM. ^#^p < 0.05 vs. D39; *p < 0.05 vs. D39 + 25-HC; n = 3–11. **(D)** Lung perfusion *in situ*. Fluorescent surface images of isolated, buffer-perfused mouse lungs infused with FITC-dextran to visualize microcirculatory perfusion (white signal, black arrows). Homogeneous perfusion was observed in control lungs (buffer only), and in lungs treated with D39Δ*ply* (5×10⁵ CFU/mL) or D39 (5×10⁵ CFU/mL) after pretreatment with 25-HC (50 µM). In contrast, lungs treated with D39 alone exhibited pronounced perfusion failure, visible as areas lacking FITC-dextran signal (black regions as negative contrast, white arrowheads). **(E)** Fluorescent surface images of a subpleural pulmonary artery with intravascular clot formation visible as areas lacking FITC-dextran signal with aggregates of non-labeled RBCs (as negative contrast) with red labeled platelets in between. **(F)** Pulmonary edema formation *in situ*. Two-photon microscopy images of subpleural pulmonary arterioles from isolated, buffer-perfused mouse lungs infused with FITC-dextran (green) to visualize vascular leakage. Leukocytes were labeled with anti-Ly6G antibody (red), and UV auto-fluorescence (blue) highlights extracellular matrix. Upper panel: xy-view. In D39-treated lungs (5×10⁵ CFU/mL, left), interstitial (arrowheads) and intra-alveolar (arrows) edema were observed, whereas infusion of HEPES alone, D39ΔPLY, D39 under Ca^2+^-free conditions, or 25-HC pretreatment (50 µM, right) prevented extravasation. Lower panel: orthogonal Z-stack view of the same arterioles, delineating vessel margins. FITC-dextran extravasation into the interstitium occurred after D39, but remained confined within the lumen after HEPES, D39ΔPLY, D39 under Ca^2+^-free conditions, or 25-HC pretreatment. **(G)** Group data of mean lung wet/dry weight ratios under baseline conditions (control), or after treatment with D39 (5×10⁵ CFU/mL), D39Δ*ply* (5×10⁵ CFU/mL), or D39 with 25-HC pretreatment (50 µM), as indicated. **(H)** Group data of mean pulmonary artery pressure (PAP) responses shown as percentage change from baseline under the same treatment conditions. Data are presented as Mean ± SEM. *p < 0.05 vs. control; ^#^p < 0.05 vs. D39; n = 4–12.

High-resolution two-photon imaging revealed that D39 were frequently retained within pulmonary microvessels and were associated with adherent leukocytes forming intravascular aggregates (Fig. 2A). Ly6G⁺ leukocytes were observed catching GFP-expressing D39 (Fig. 2B) and forming extracellular DNA structures consistent with neutrophil extracellular traps (NETs) ((Fig. 2C, D). In addition, fine filamentous connections resembling tunneling nanotubes (TNTs) were occasionally observed between adjacent leukocytes (Fig. 2E) and bacteria were often detected within alveolar capillary loops, consistent with mechanical trapping at the microvascular interface (Fig. 2E). These observations indicate that pneumococcal infection rapidly induces pulmonary vascular injury accompanied by intravascular cell recruitment and bacterial trapping.

**Figure 2.**
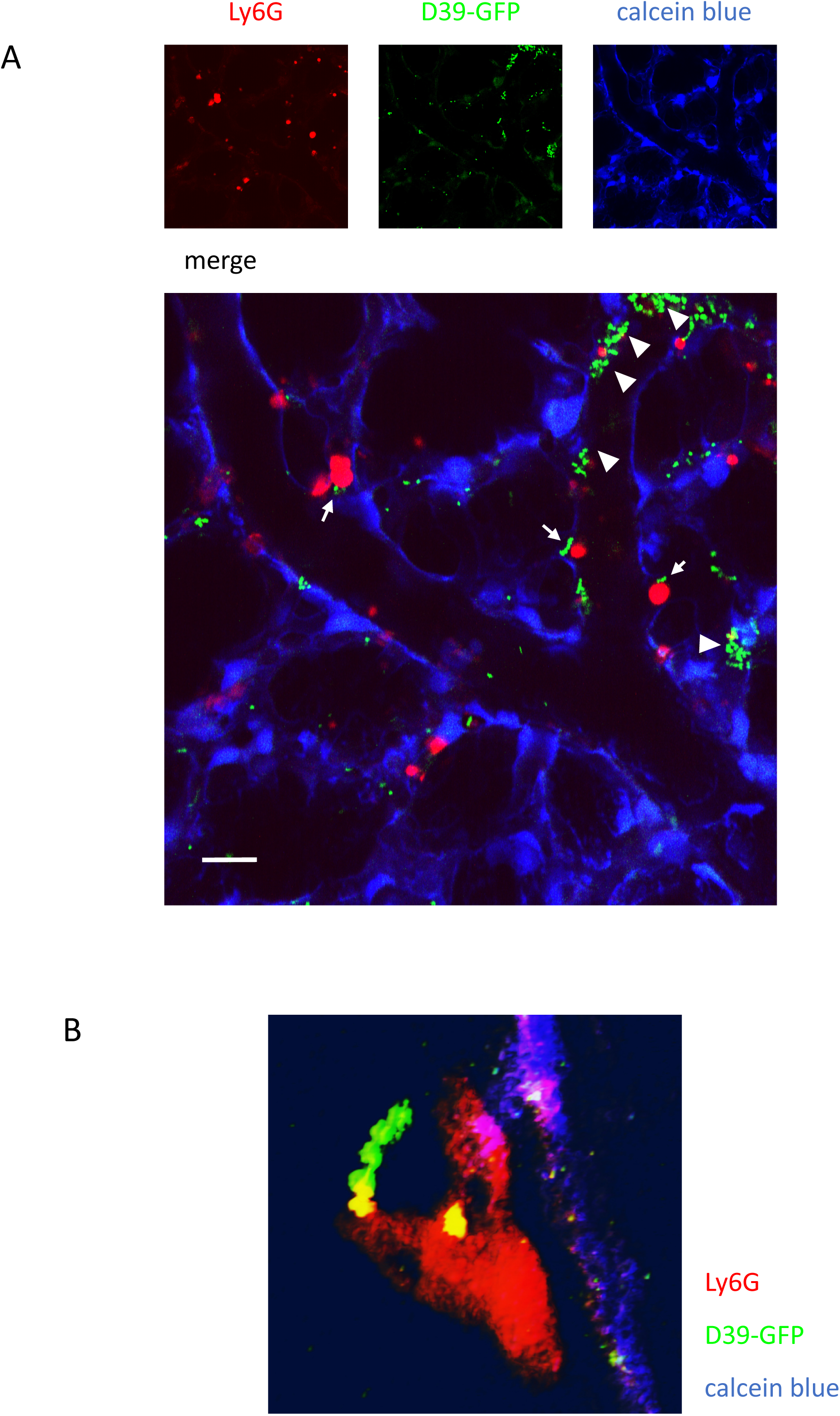
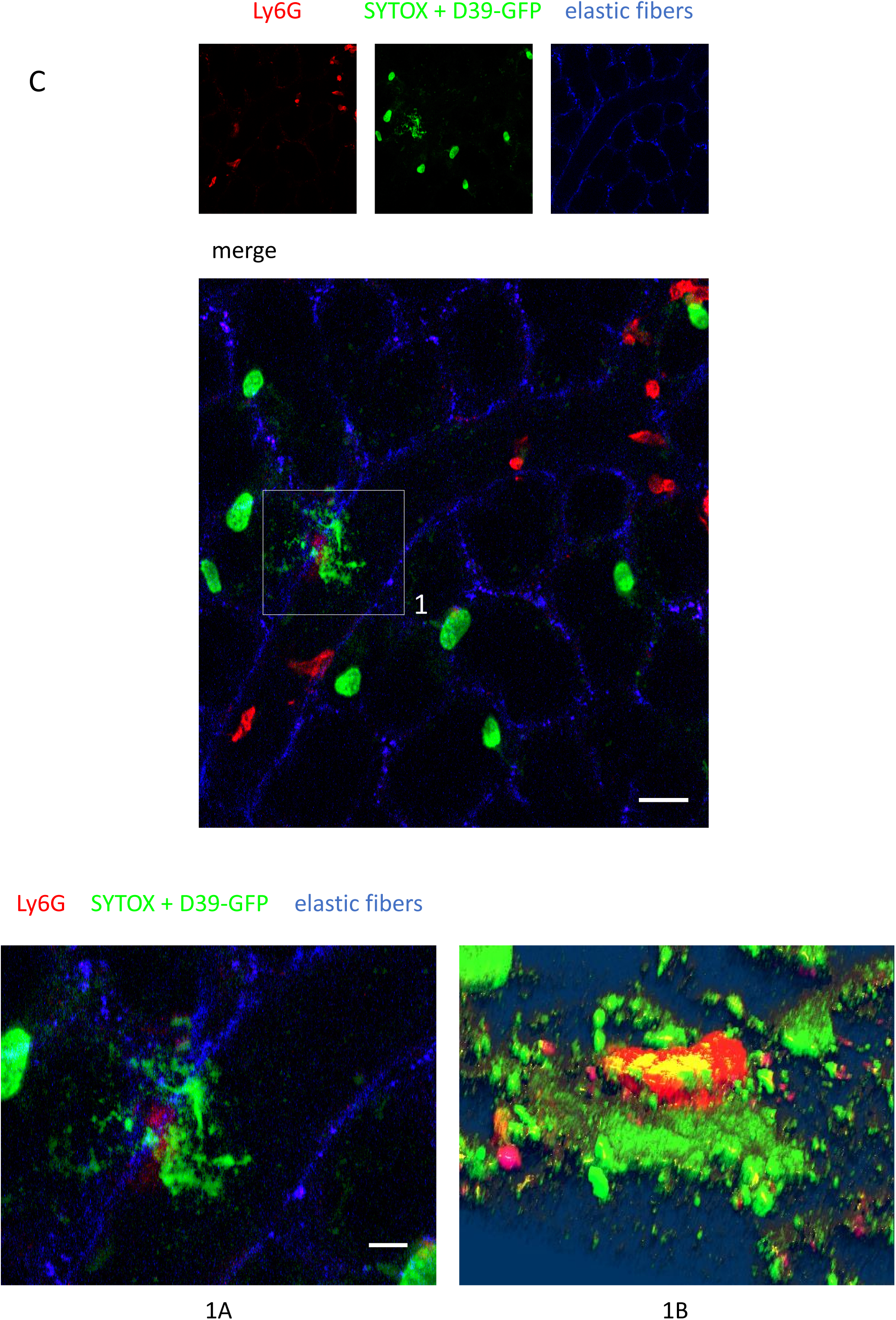
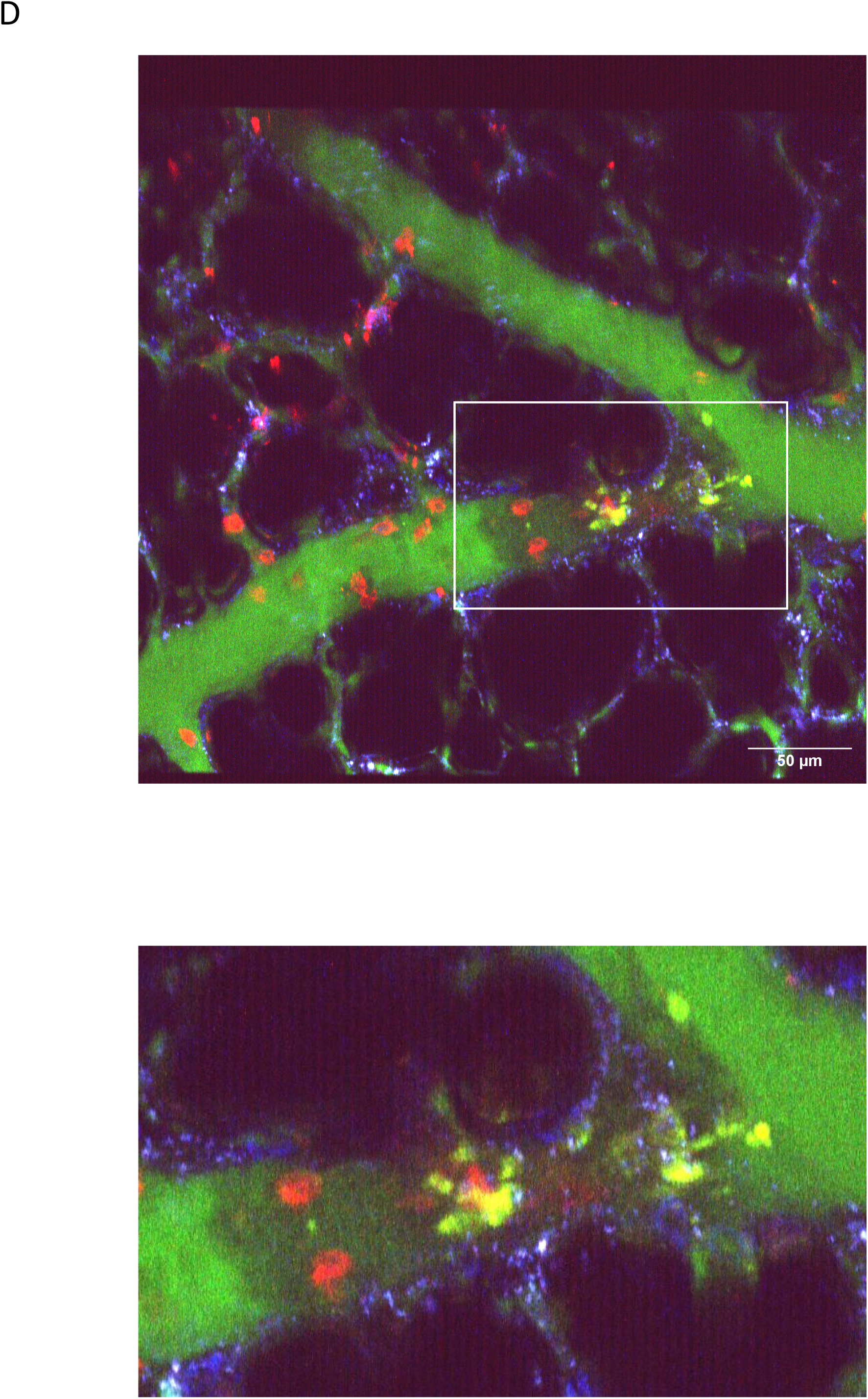
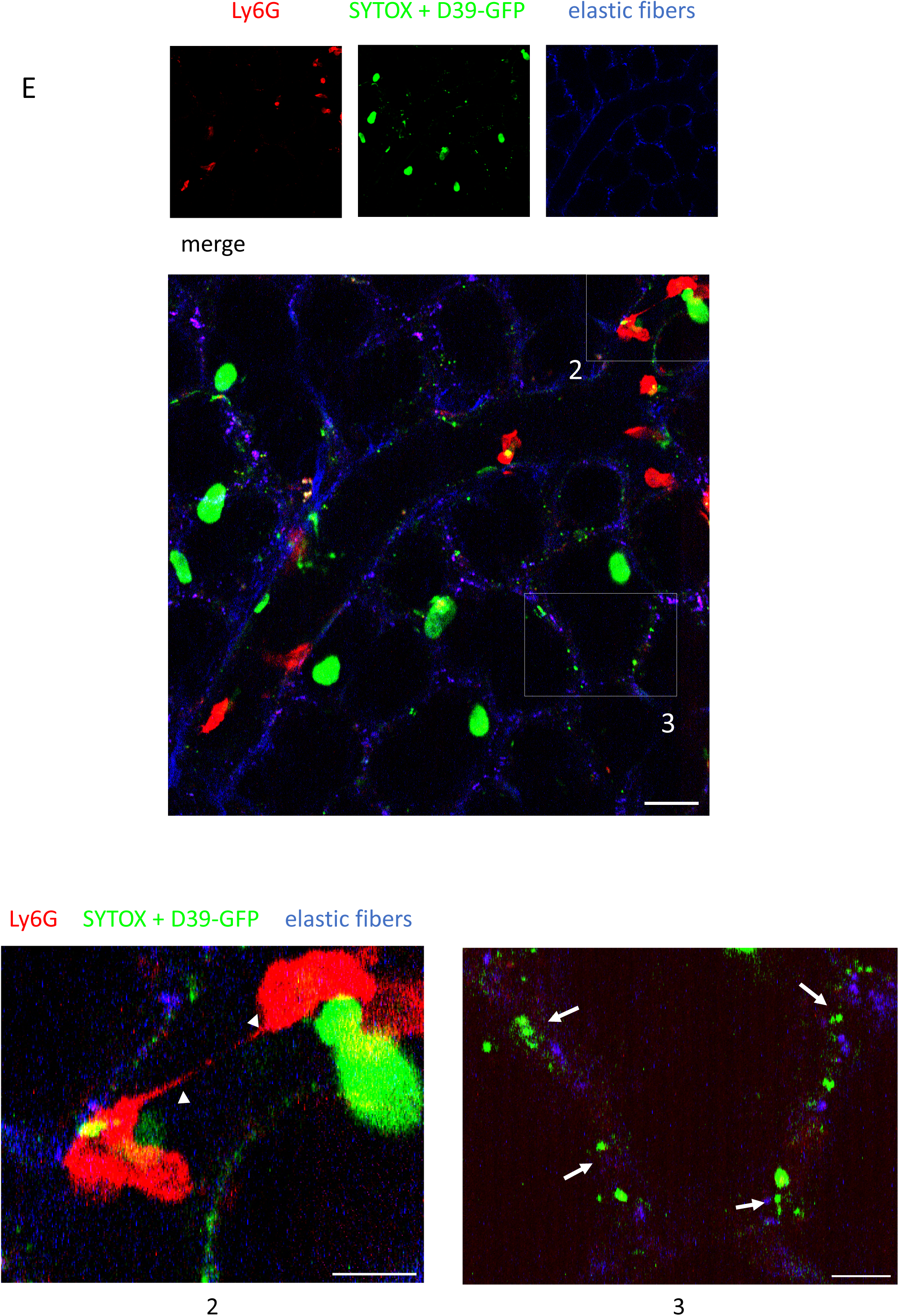
**(A)** Representative two-photon image of a subpleural pulmonary arteriole in an isolated, buffer-perfused mouse lung following intravascular infusion of GFP-expressing D39. Ly6G⁺ leukocytes are shown in red and GFP-labeled bacteria in green. Endothelium is stained with calcein blue. The lower panel depicts the merged image; upper panels display the individual fluorescence channels separately. Scale bar, 25 µm. In the merged image **arrows heads** indicate GFP-positive *D39* aggregates and **arrows** indicate trapping of GFP-positive *D39* by a Ly6G⁺ cell; **(B)** D39 trapping is further shown at higher magnification in the corresponding subpanels. **(C)** Two-photon image of a subpleural pulmonary arteriole in an isolated, buffer-perfused mouse lung following intravascular infusion of WT D39. Ly6G⁺ leukocytes are shown in red, SYTOX in green, and tissue autofluorescence of elastic fibers in blue. The lower panel depicts the merged image; upper panels display the individual fluorescence channels separately. Scale bar, 25 µm. In the merged image, SYTOX fluorescence indicating NET formation surrounding red labeled leukocyte; framed region 1 is shown at higher magnification in the corresponding subpanels in 2D (left) and as 3D (right) reconstruction, respectively. **(D)** Two-photon image of a subpleural pulmonary arteriole in an isolated, buffer-perfused mouse lung following intravascular infusion of WT D39. Ly6G⁺ leukocytes are shown in red, FITC dextrane in green, SYTOX in yellow, and tissue autofluorescence of elastic fibers in blue. Scale bar, 25 µm. In the merged image, SYTOX fluorescence indicating NET formation surrounding red labeled leukocyte in the center of a bulk of RBCs (as negative contrast) indicting clot formation. Vessel perfusion seemed to be maintained by retrograde collaterale inward currents. Framed region 1 is shown at higher magnification in the corresponding subpanels **(E)** Two-photon image of a subpleural pulmonary arteriole in an isolated, buffer-perfused mouse lung following intravascular infusion of GFP-expressing D39. Ly6G⁺ leukocytes are shown in red, GFP-labeled bacteria and SYTOX uptake in green, and tissue autofluorescence of elastic fibers in blue. The lower panel depicts the merged image; upper panels display the individual fluorescence channels separately. Scale bar, 25 µm. In the merged image, representative features are highlighted: frame 2, tunneling nanotube-like (TNT) connections between adjacent Ly6G⁺ leukocytes; frame 3, GFP-labeled pneumococci trapped within alveolar capillaries. Each framed region is shown at higher magnification in the corresponding subpanels. Arrows indicate TNT and S. pneumoniae, respectively.

Because PLY is a major virulence factor of *S. pneumoniae*, we next asked whether the observed microvascular injury depends on toxin expression. Lungs perfused with a PLY-deficient strain (D39Δply) showed minimal platelet and leukocyte recruitment, no increase in pulmonary artery pressure or wet-to-dry weight ratio, and only minor impairment of microvascular perfusion compared with wild-type bacteria indicating that PLY is required for both vascular injury and intravascular trapping (Fig. 1A–H).

PLY is known to form PM pores that allow influx of extracellular Ca²⁺ ^16,17^. We therefore considered that Ca²⁺ entry may be required for the vascular responses observed after pneumococcal exposure. To test this hypothesis, lungs were perfused with Ca²⁺-free buffer during D39 infusion. Under Ca²⁺-free conditions, platelet adhesion and leukocyte recruitment were strongly reduced, and D39 failed to induce significant perfusion failure, pulmonary edema, or vascular leakage (Fig. 1A–H). These findings suggest that pneumococcal lung injury depends on a Ca²⁺-dependent process consistent with toxin-induced PM permeabilization.

Because PLY PM pore formation requires accessible PM cholesterol ^7^, we next examined whether modification of PM lipid composition affects the development of lung injury. Pretreatment of lungs with 25-HC, an oxysterol known to reduce accessible PM cholesterol, markedly attenuated D39-induced leukocyte and platelet recruitment, preserved microvascular perfusion, and reduced pulmonary edema and vascular leakage, suggesting that inhibition of PLY activity protects against vascular injury but may simultaneously limit host responses associated with bacterial trapping (Fig. 1A–H).

Together, these findings indicate that pneumococcal lung injury requires PLY and extracellular Ca²⁺ and can be prevented by the cholesterol-modifying lipid 25-HC, consistent with a mechanism involving toxin-induced PM pore formation.

### Pneumolysin-dependent membrane permeabilization and Ca²⁺ signaling in pulmonary microvessels are inhibited by 25-hydroxycholesterol

The experiments shown in Fig. 1 indicated that pneumococcal lung injury requires PLY, depends on extracellular Ca²⁺, and is prevented by 25-HC, suggesting that toxin-induced PM pore formation may trigger Ca²⁺ influx in the pulmonary microvasculature. To test this hypothesis directly, we examined PM permeabilization and intracellular Ca²⁺ signaling in pulmonary vessels *in situ*.

To visualize PM damage in the intact lung, we monitored uptake of the PM-impermeable nucleic acid dye SYTOX during perfusion of isolated lungs with GFP-expressing D39. In control lungs, SYTOX fluorescence was absent, indicating intact endothelial PMs (data not shown). In contrast, exposure to wild-type D39 caused rapid SYTOX uptake in endothelial cells of subpleural arterioles, consistent with toxin-induced PM permeabilization (Fig. 3A).

**Figure 3.**
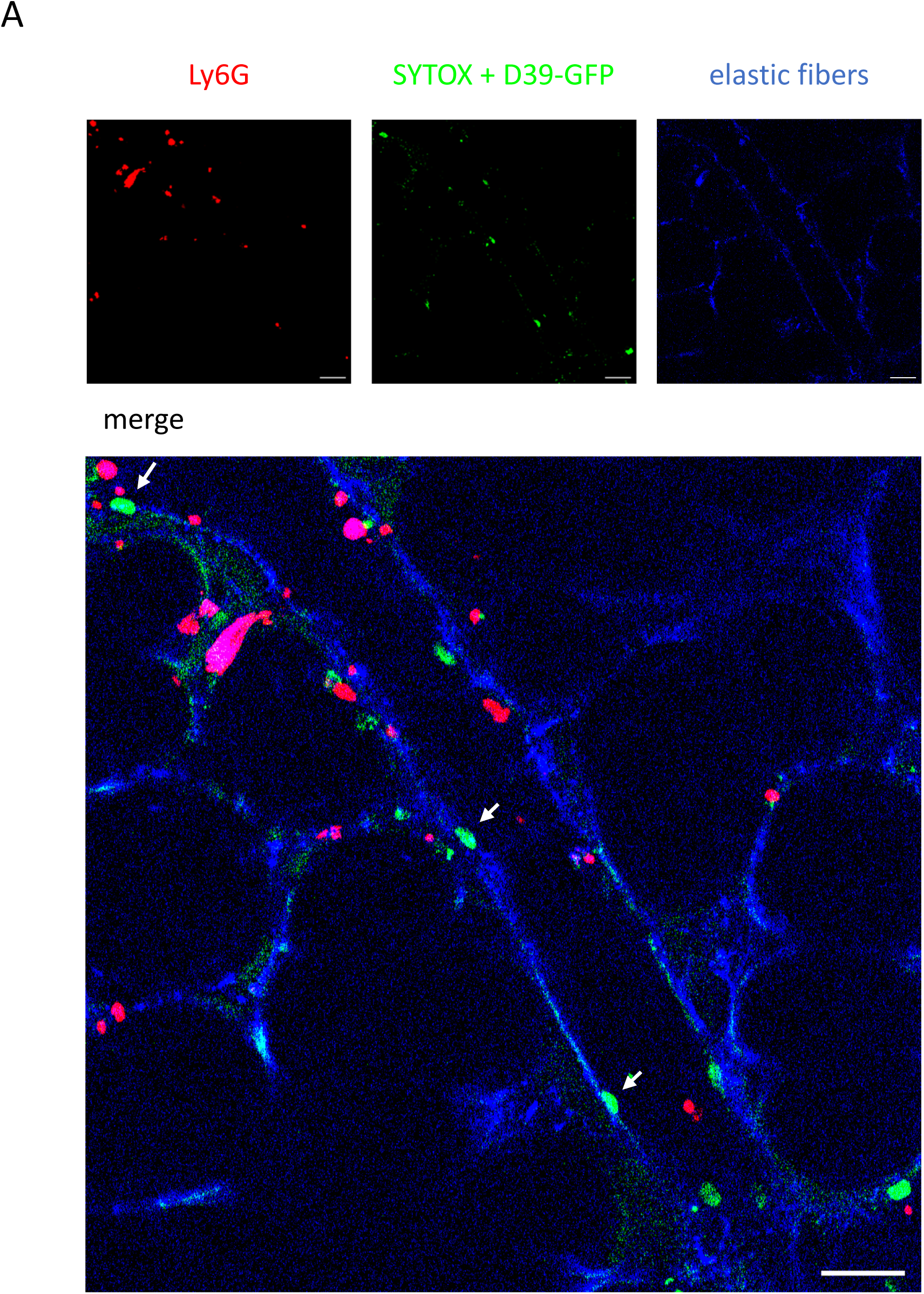
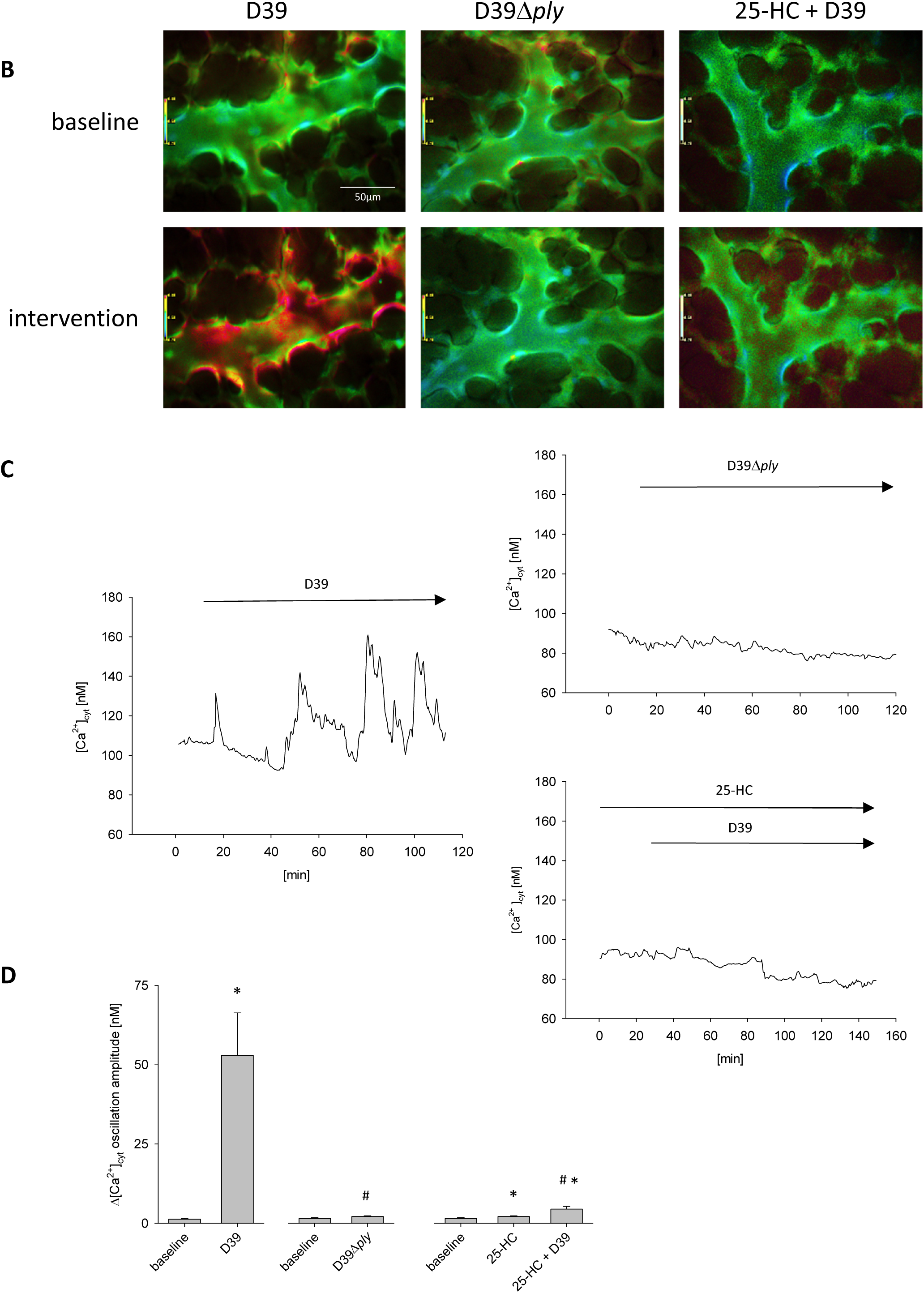
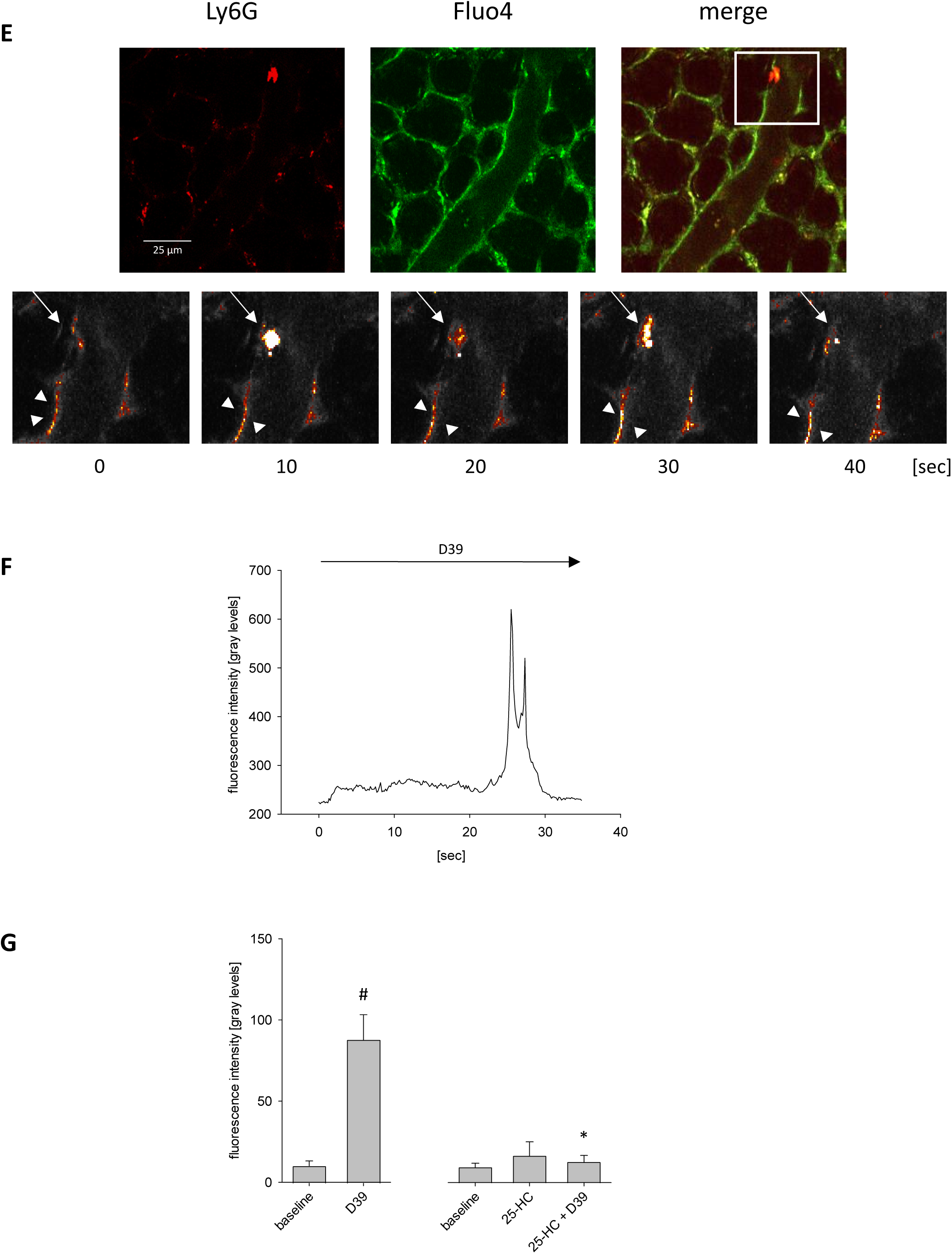
PLY-induced PM pore formation and Ca²⁺ signaling in pulmonary arteriolar endothelial cells and leukocytes *in situ*. **(A)** Representative two-photon image of a subpleural pulmonary arteriole in an isolated, buffer-perfused mouse lung following intravascular infusion of WT D39. Ly6G⁺ leukocytes are shown in red, SYTOX in green, and tissue autofluorescence of elastic fibers in blue. The lower panel depicts the merged image; upper panels display the individual fluorescence channels separately. Scale bar, 25 µm. In the merged image, SYTOX uptake in endothelial cells (arrows) indicating plasma-membrane permeabilization. **(B)** Pseudocolor-coded images of the 340:380 ratio in fura-2–loaded endothelial cells lining subpleural arterioles of isolated, buffer-perfused mouse lungs. Recordings were taken under baseline conditions and after intravascular application of D39 (5×10⁵ CFU/mL), D39Δ*ply* (5×10⁵ CFU/mL), or D39 following pretreatment with 25-HC (50 µM), as indicated. **(C)** Representative [Ca²⁺]_cyt_ tracings from arteriolar endothelial cells before and after treatment with D39, D39Δ*ply*, or D39 + 25-HC. **(D)** Group data showing mean Ca²⁺ oscillation amplitude within a 10-minute baseline and a 30-minute post-treatment interval. Data are presented as Mean ± SEM. *p < 0.05 vs. baseline; ^#^p < 0.05 vs. D39; n = 6–12. **(E)** Top row: Two-photon microscopy images of a subpleural arteriole from an isolated, buffer-perfused mouse lung after D39 (5×10⁵ CFU/mL) infusion. Left: Ly6G-labeled adherent leukocytes (red). Middle: Fluo-4 fluorescence in leukocytes and endothelial cells (green). Right: Merged image. Bottom row: High-magnification time series (10-second intervals) of the region marked in the merged image. Fluo-4 fluorescence in the adherent leukocyte (arrow) and adjacent endothelial cells (arrowheads) demonstrates Ca²⁺ oscillations in both cell types. **(F)** Representative [Ca²⁺]_cyt_ tracing from the marked leukocyte after D39 treatment. **(G)** Group data of maximal [Ca²⁺]_cyt_ response in leukocytes under baseline conditions and within 30 minutes following D39 (5×10⁵ CFU/mL), 25-HC (50 µM), or their combination. Data are shown as Mean ± SEM. ^#^p < 0.05 vs. baseline; *p < 0.05 vs. D39; n = 6–12.

We next asked whether PLY-induced PM damage is associated with Ca²⁺ influx in the pulmonary microvasculature. Using fura-2–based imaging in isolated perfused lungs, we observed that application of wild-type D39 induced repetitive increases in intracellular Ca²⁺ in endothelial cells of subpleural arterioles (Fig. 3B-D). These Ca²⁺ signals were not detected in lungs exposed to D39Δply, indicating that Ca²⁺ entry requires toxin expression (Fig. 3B-D). Importantly, pretreatment with 25-HC strongly attenuated the Ca²⁺ response, consistent with inhibition of PM pore formation (Fig. 3B-D).

Because leukocyte recruitment was prominent in Fig. 1, we also examined Ca²⁺ signaling in intravascular leukocytes. In LY6G-labeled, Fluo-4–loaded lungs, exposure to wild-type D39 triggered transient Ca²⁺ spikes in adherent leukocytes (Fig. 3E-G), whereas the Δply mutant failed to evoke detectable signals (not shown). Similar to endothelial cells, leukocyte Ca²⁺ responses were markedly reduced after treatment with 25-HC (Fig. 3G).

Together, these findings demonstrate that PLY induces PM permeabilization and Ca²⁺ influx in endothelial cells and leukocytes within the pulmonary microvasculature and that both responses are inhibited by 25-HC. These results support the hypothesis that toxin-induced PM pore formation triggers the Ca²⁺-dependent vascular injury observed in Fig. 1.

### Pneumolysin forms membrane pores that mediate extracellular Ca²⁺ entry

The *in situ* experiments demonstrated PLY-dependent PM permeabilization and Ca²⁺ signaling in pulmonary microvessels and suggested that toxin-induced PM pore formation allows influx of extracellular Ca²⁺. To directly test this mechanism, we analyzed PLY-induced PM permeabilization and Ca²⁺ entry in isolated cells *in vitro*.

As a classical readout for PM pore formation by CDCs, hemolysis of erythrocytes was used as a sensitive indicator of PM damage ^18,19^. Both recombinant PLY and wild-type D39 induced concentration-dependent hemolysis, whereas the PLY-deficient mutant had no effect, confirming that PM pore formation depends on PLY expression (Supplementary Fig. S1).

To determine whether PLY permeabilizes pulmonary endothelial cells, PM integrity was assessed in human pulmonary microvascular endothelial cells (HPMECs) using the PM-impermeable dye SYTOX green. Exposure to D39 or recombinant PLY caused progressive intracellular accumulation of SYTOX fluorescence, indicating loss of PM integrity, whereas the D39Δply failed to induce detectable uptake (Fig. 4A, B). As expected, Triton X-100, used as a positive control, also caused a marked increase in SYTOX uptake (fig. 4B). These findings demonstrate that PLY directly permeabilizes endothelial cell PMs. Consistent with both, the RBC hemolysis and the *in situ* results, 25-HC pretreatment completely inhibited SYTOX uptake following D39 or PLY exposure (Figure 4A, B). Importantly, 25-HC did not prevent SYTOX entry in Triton X-treated cells, confirming that its protective effect is specific to PLY-mediated PM pore formation and not a general inhibitor of PM permeability (Figure 4B).

**Figure 4.**
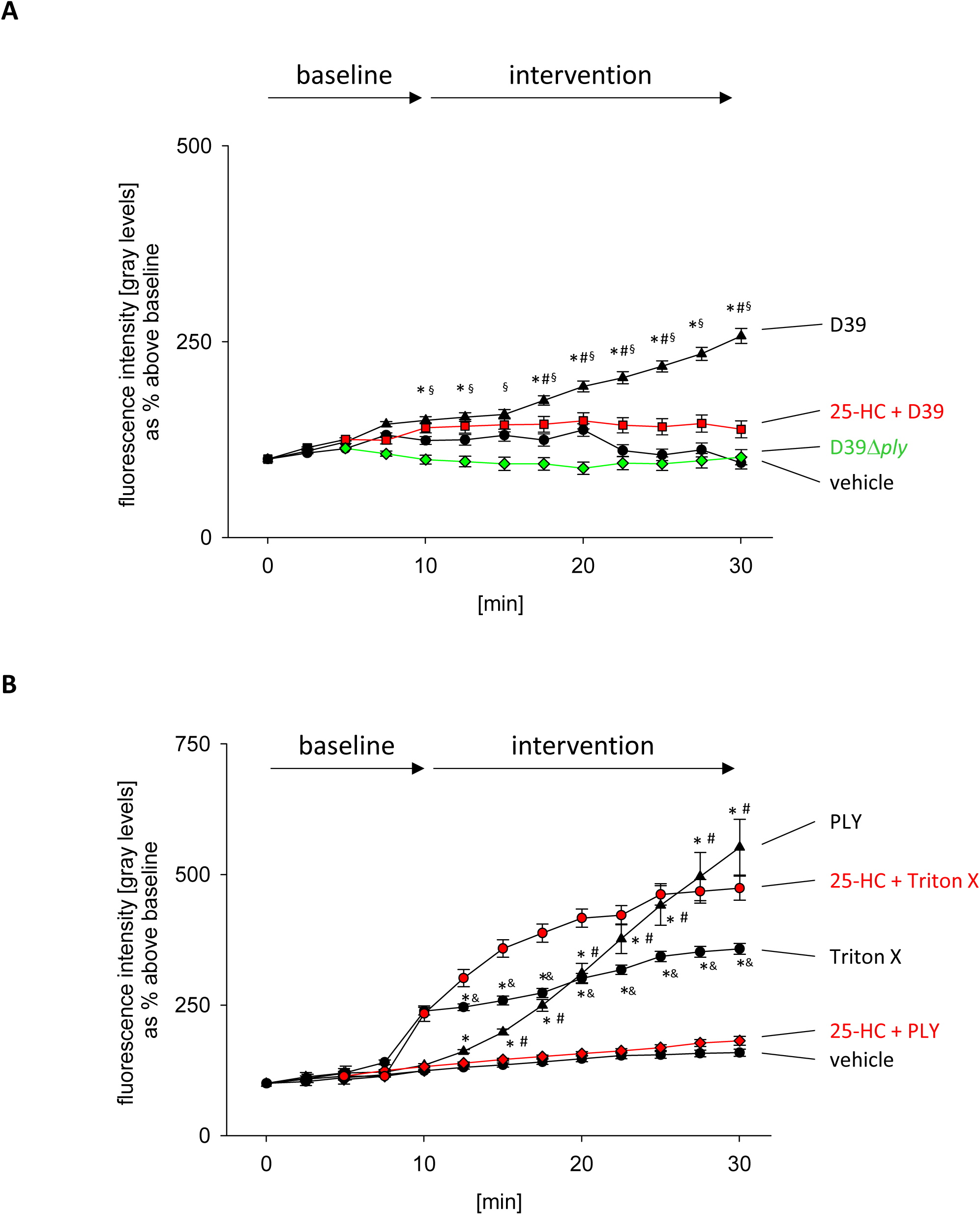
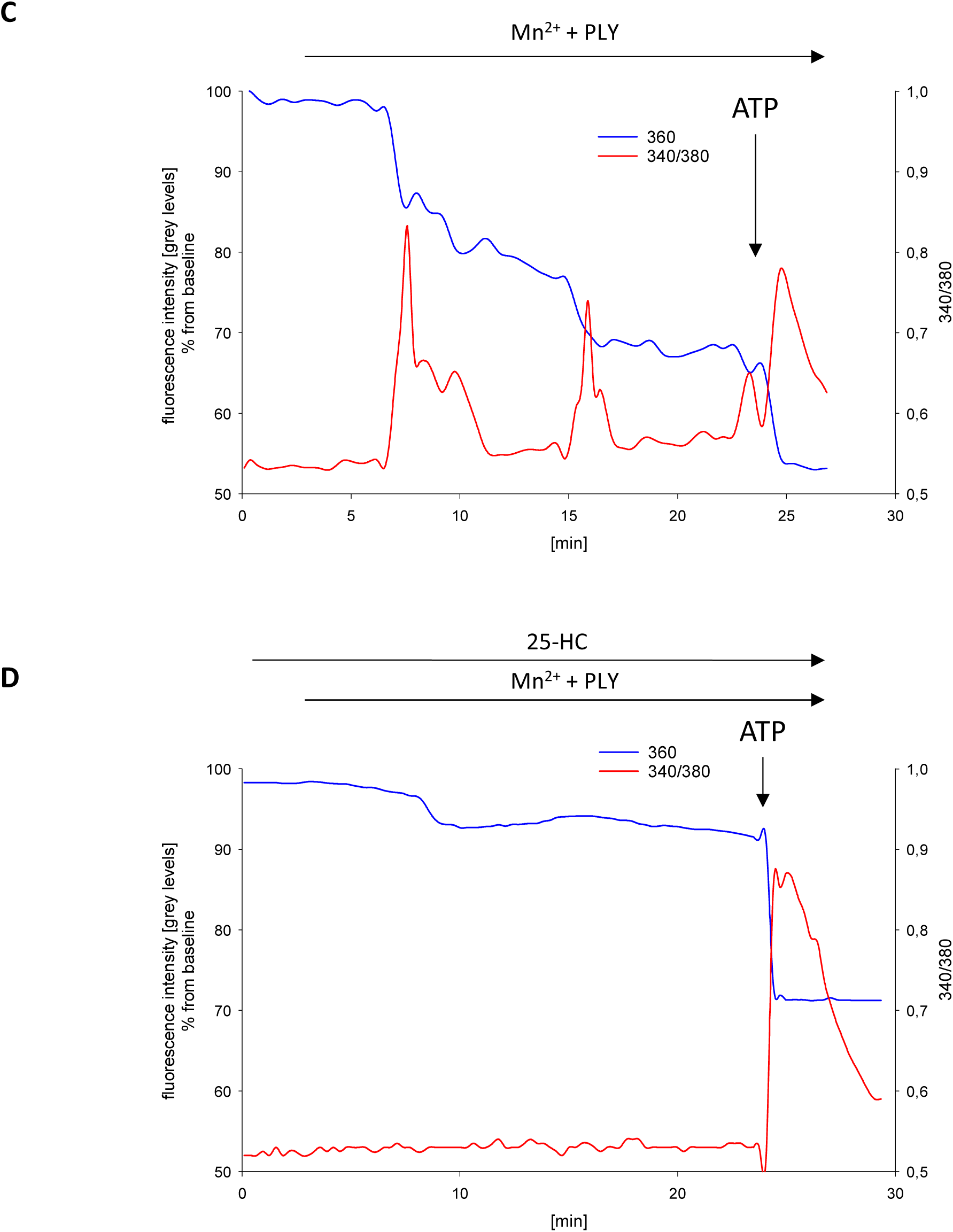
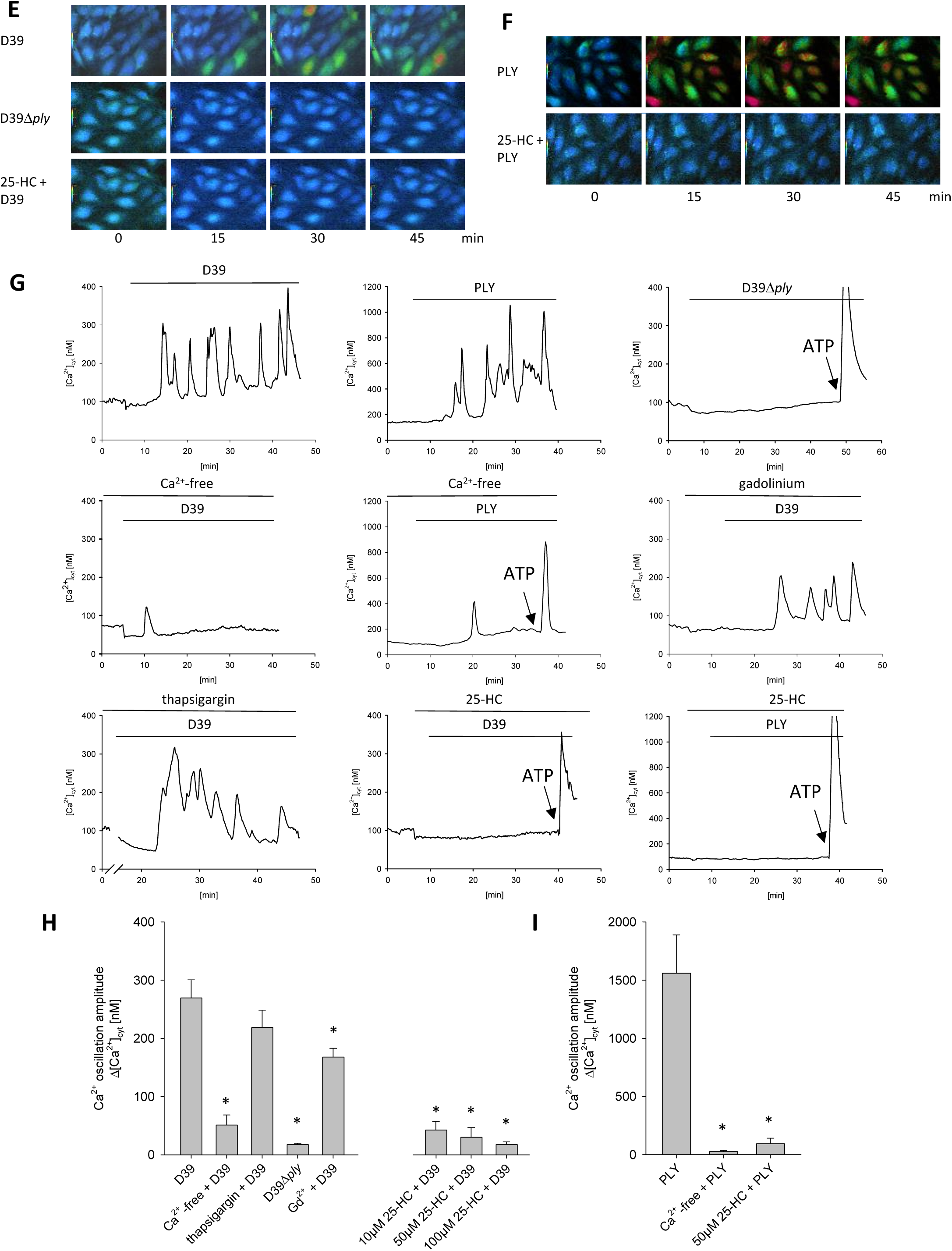
PLY-induced PM pore formation and Ca²⁺ signaling in HEPMECs *in vitro*. **(A, B)** Plasma membrane pore formation in HPMECs assessed by uptake of SYTOX, a membrane-impermeant DNA dye. Group data showing SYTOX fluorescence as percent of baseline values before and after exposure to D39 (5×10⁵ CFU/mL), D39Δ*ply* (5×10⁵ CFU/mL), PLY (50 ng/mL), vehicle as negative control, or Triton X-100 (0.01%) as positive control, without or with 25-HC pretreatment, as indicated. 25-HC prevented SYTOX uptake in response to PLY and D39 but not Triton X. Data are presented as mean ± SEM; ^#^p < 0.05 vs. vehicle, *p < 0.05 vs. 25-HC, ^§^p < 0.05 vs. D39Δ*ply*, ^&^p < 0.05 vs. Triton X + 25-HC; n = 4–9. **(C,D)** PLY-mediated pore formation induces external Ca²⁺ influx in HPMECs, which is blocked by 25-HC. Representative traces of fura-2 fluorescence ratio (340/380) and corresponding 360 nm fluorescence in HPMECs loaded with fura-2 AM and exposed to Ca²⁺-containing buffer. After a brief baseline, Mn²⁺ was added to initiate quenching of fura-2 at 360nm, followed by treatment with PLY (50ng/ml) alone (C) or in combination with 25-HC (50 µM) (D). ATP (100 µM) was applied as a positive control at the end of each experiment. PLY induced Mn²⁺ quenching and Ca²⁺ influx, indicative of pore formation, which was abolished by 25-HC.PLY-mediated pore formation induces external Ca²⁺ influx in HPMECs, which is blocked by 25-HC. Representative traces of fura-2 fluorescence ratio (340/380) and corresponding 360 nm fluorescence in HPMECs loaded with fura-2 AM and exposed to Ca²⁺-containing buffer. After a brief baseline, Mn²⁺ was added to initiate quenching of fura-2 at 360nm, followed by treatment with PLY (50ng/ml) alone (C) or in combination with 25-HC (50 µM) (D). ATP (100 µM) was applied as a positive control at the end of each experiment. PLY induced Mn²⁺ quenching and Ca²⁺ influx, indicative of pore formation, which was abolished by 25-HC. **(E, F)** Time-lapse pseudocolor images of fura-2 340/380 ratio in HPMECs exposed to D39 (5×10⁵ CFU/mL), D39Δ*ply* (5×10⁵ CFU/mL), or purified PLY (50 ng/mL), with or without 25-HC (50 µM), as indicated. **(G)** Representative [Ca²⁺]_cyt_ traces showing effects of extracellular Ca²⁺ removal or pharmacological pretreatment with thapsigargin (10 µM), gadolinium (1 mM), or 25-HC on responses to D39, D39Δ*ply*, or PLY, as indicated. ATP (100 µM) was added at the end as a positive control. **(H, I)** Grouped data quantifying mean Ca²⁺ oscillation amplitude in HPMECs under indicated conditions. Data are Mean ± SEM. *p < 0.05 vs. D39 or PLY; n = 5–26.

For direct proof that PLY-induced PM pores enable Ca²⁺ entry into HPMECs, we next used manganese (Mn²⁺) quenching as a surrogate marker of divalent cation influx. Fura-2–loaded HPMECs were exposed to extracellular Mn²⁺, and fluorescence was recorded at 360 nm excitation, which is the isosbestic wavelength where fura-2 fluorescence is independent of Ca²⁺ levels. A decline in 360 nm fluorescence indicates Mn²⁺ entry across a compromised PM ^20^. Upon exposure to PLY, we observed a delayed onset (5–10 min) of transient [Ca²⁺]_cyt_ spikes accompanied by Mn²⁺ quenching of 360 nm fluorescence. Notably, the Mn²⁺ signal plateaued after an initial drop, suggesting that pore-mediated influx was rapidly terminated, presumably due to PM resealing. These observations support the interpretation that PLY forms transient PM pores that allow Ca²⁺ and Mn²⁺ influx (Figure 4C).

Pre-treatment with 25-HC completely abolished both Mn²⁺ quenching and PLY-induced Ca²⁺ transients, consistent with its inhibitory effect on PM pore formation (Fig. 4D). To test whether 25-HC also affects classical Ca²⁺ channel activity, we stimulated HPMECs with ATP, a known activator of both intracellular Ca²⁺ release from endoplasmic reticulum (ER) and store-operated Ca²⁺ entry (SOCE). ATP induced a characteristic [Ca²⁺]_cyt_ spike along with Mn²⁺ influx, neither of which were altered by 25-HC, indicating that SOCE channels remain functional in the presence of 25-HC (Fig. 4C, D). These results suggest that 25-HC does not interfere with canonical Ca²⁺ channels but specifically inhibits PM pore-mediated Ca²⁺ entry.

We further investigated PLY-dependent Ca²⁺ signaling in fura-2–loaded HPMECs in the absence of extracellular Mn²⁺ (Fig. 3A–E). Both D39 and recombinant PLY elicited repetitive Ca²⁺ oscillations, whereas the PLY-deficient mutant D39Δ*ply* did not trigger any Ca²⁺ signal (Fig. E-I). Notably, these oscillations were abolished under Ca²⁺-free extracellular conditions, confirming their dependence on extracellular Ca²⁺ influx (Fig. 4G-I).

To explore the involvement of alternative Ca²⁺ sources, we applied the non-specific Ca²⁺ channel blocker gadolinium and the sarco/endoplasmic reticulum Ca²⁺ ATPase (SERCA) inhibitor thapsigargin. Neither treatment affected D39-induced Ca²⁺ oscillations, further supporting a mechanism reliant on PM pore-mediated Ca²⁺ entry rather than PM channel activation or ER store release (Fig. G-H). Finally, 25-HC inhibited both D39- and PLY-induced Ca²⁺ oscillations in a concentration-dependent manner (Fig. 4E-I).

Together, these experiments demonstrate that PLY forms PM pores that allow influx of extracellular Ca²⁺ into endothelial cells and provide a direct mechanistic explanation for the Ca²⁺-dependent pulmonary vascular injury observed in Fig. 1 and Fig. 2. This process is not mediated by classical Ca²⁺ channels or ER Ca²⁺ release and is potently inhibited by 25-HC, highlighting its specific and effective blockade of PLY-induced PM disruption.

### 25-Hydroxycholesterol inhibits pneumolysin pore formation by reducing accessible membrane cholesterol

PLY, a CDC, requires accessible PM cholesterol to bind and form pores in the PM. 25-HC, an oxysterol, has been shown to regulate cellular cholesterol levels. To elucidate the mechanism by which 25-HC inhibits PLY activity, we focused on its known ability to activate ACAT, thereby promoting esterification of free cholesterol and reducing its PM accessibility ^7–9^.

Consistent with this mechanism, pharmacological inhibition of ACAT using Sandoz 58-035 reversed the protective effect of 25-HC, restoring both PLY-induced PM pore formation and cytosolic Ca²⁺ signaling in HPMECs (Figure 5A, B). To further confirm that 25-HC exerts its effect via cholesterol accessibility, we pretreated HPMECs with sphingomyelinase (SMase), which hydrolyzes sphingomyelin to ChoP and ceramide. Because a significant proportion of PM cholesterol is sequestered in sphingomyelin-rich domains and thereby inaccessible to CDCs, SMase treatment increases the pool of accessible cholesterol ^7^. Indeed, SMase pretreatment also abolished the inhibitory effect of 25-HC on PLY-induced Ca²⁺ signaling, further supporting the cholesterol accessibility hypothesis (Figure 5A, B).

**Figure 5.**
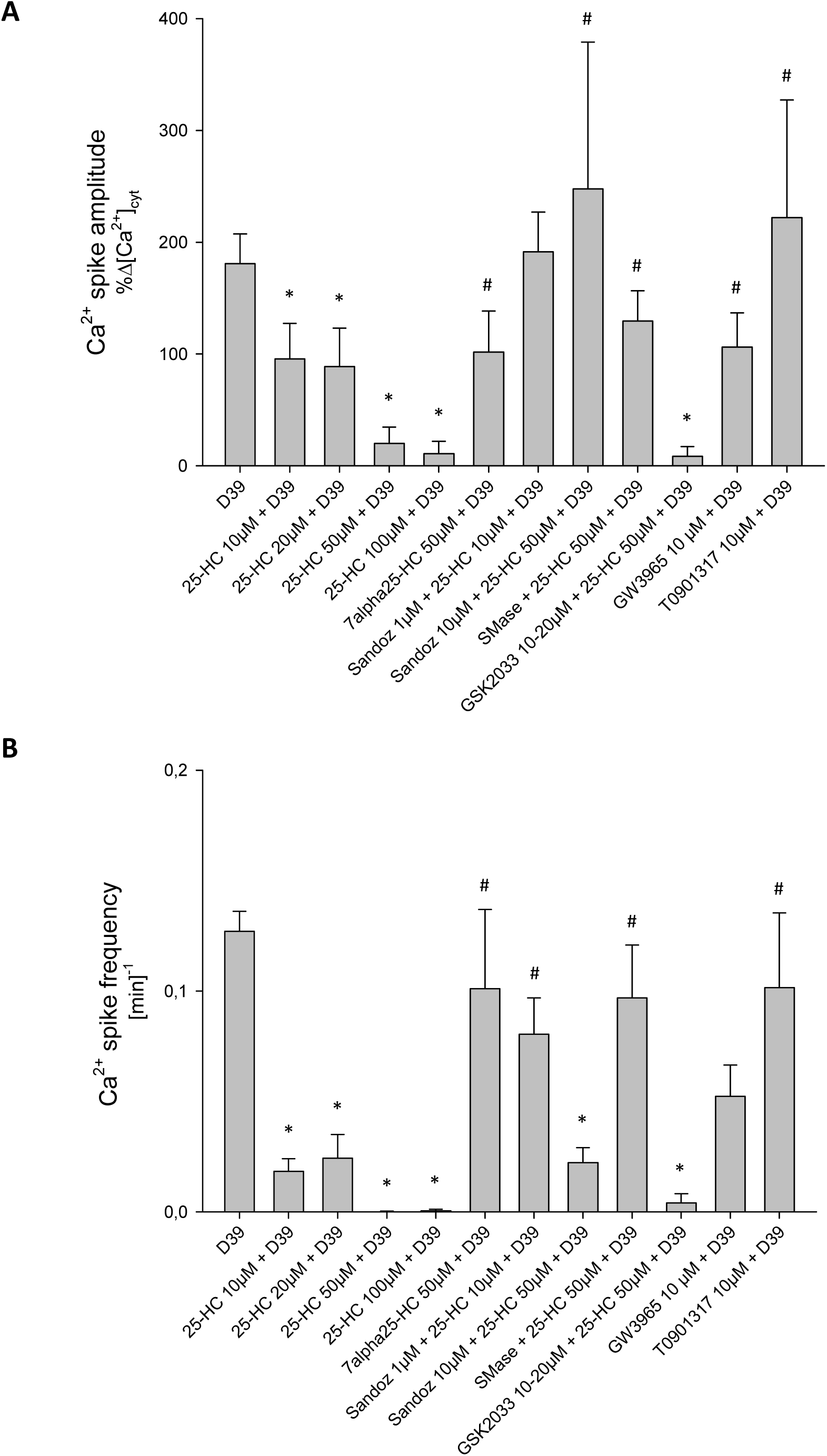
Role of ACAT in 25-HC-mediated inhibition of D39-induced Ca²⁺ oscillations in HPMECs. **(A)** Grouped data showing mean Ca²⁺ oscillation amplitudes as percentage of baseline over a 30-minute recording period. **(B)** Corresponding spike frequencies under the same conditions. HPMECs were treated with D39 (5×10⁵ CFU/mL) alone or in combination with 25-HC (10-100 µM), 7α,25-HC (50 µM), Sandoz 58-035 (1 µM), SMase (50 mU/mL), or LXR modulators GSK2033 (20µM), GW3965 (10µM), or TO191317 (10 µM), as indicated. Data are Mean ± SEM. *p < 0.05 vs. D39 alone, ^#^p < 0.05 vs. 25-HC (50 µM) + D39; n = 4–29.

In addition to activating ACAT, 25-HC has been reported to serve as an agonist of liver X receptors (LXRs), nuclear receptors involved in lipid metabolism ^21–23^. To assess the potential involvement of LXRs in 25-HC’s action, we pretreated HPMECs with the LXR antagonist GSK2033 or with LXR agonists GW3965 and T0901317. Neither the antagonist reversed the 25-HC effect, nor did the agonists mimic it. These findings exclude a major role of LXR signaling in 25-HC-mediated inhibition of PLY activity (Figure 5A, B).

Given that oxysterols can also bind estrogen receptors (ERs) ^24^, we examined whether ERs signaling contributed to the observed effects. However, treatment with the ERs agonist 7α-25-HC had no influence on PLY-induced Ca²⁺ signaling, ruling out ERs involvement (Figure 5A, B).

Together, these results demonstrate that 25-HC inhibits PLY PM pore formation by activating ACAT and reducing the pool of accessible PM cholesterol required for toxin binding and pore formation.

### Pneumococcal infection suppresses CH25H expression via inhibition of PAF receptor signaling

Because 25-HC protected against PLY-induced vascular injury, we next asked whether the CH25H–25-HC pathway is regulated during pneumococcal infection. Induction of CH25H would be expected to limit toxin-mediated damage, and therefore we initially hypothesized that pneumococcal exposure increases CH25H expression. Furthermore, 25-HC production is strongly induced during viral infections - including SARS-CoV-2 - and known to inhibit viral uptake, we thus investigated whether CH25H is also upregulated in response to *S. pneumoniae* ^6,25,26^. Surprisingly, instead of an upregulation, D39 significantly suppressed both CH25H transcription and protein expression in HPMECs (Figure 6A–B). In contrast, stimulation with IFN-γ markedly increased CH25H mRNA, and lipopolysaccharide (LPS) induced a modest upregulation, consistent with previous findings ^6,27^.

**Figure 6.**
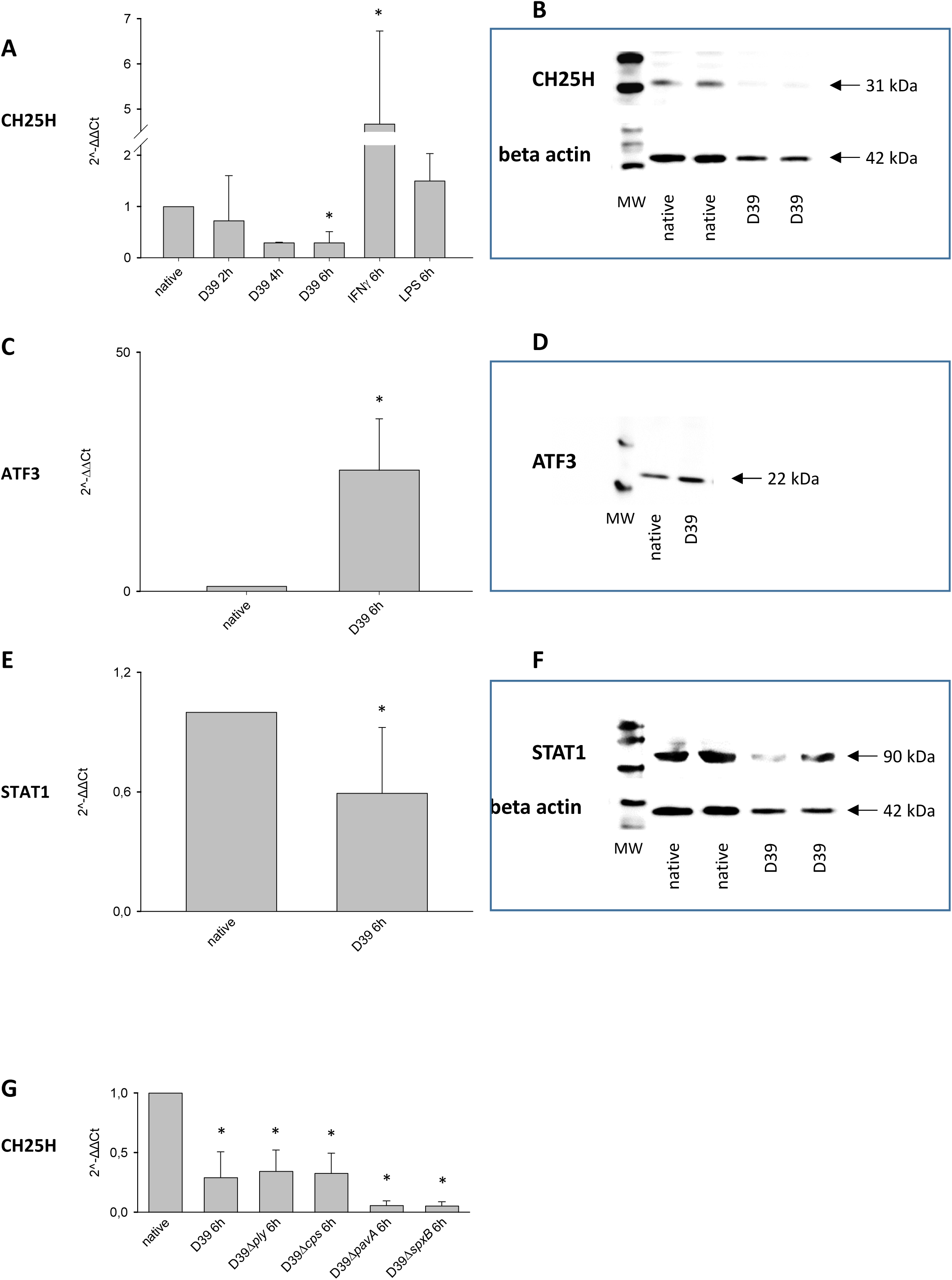
CH25H, ATF3, and STAT1 expression in HPMECs. **(A)** Group data of relative CH25H mRNA expression after 2, 4, or 6 h treatment with D39 (5×10⁵ CFU/mL), or 6 h stimulation with IFNγ (100 U/mL) or LPS (1 µg/mL), as indicated. Mean ± SEM, *p < 0.05% vs. native, n=3-4. **(B)** Representative Western blot of CH25H and β-actin protein levels in untreated (native) cells and in cells treated with D39 (5×10⁵ CFU/mL) for 6 h. **(C)** Group data of relative ATF3 mRNA expression in native cells and after 6 h D39 treatment. Mean ± SEM, *p < 0.05 vs. native, n = 3–5. **(D)** Representative Western blot of ATF3 and β-actin protein levels in native and D39-treated cells. **(E)** Representative Western blot of STAT1 and β-actin protein expression in native and D39-treated cells. **(F)** Group data of relative STAT1 mRNA expression in native and D39-treated cells. Mean ± SEM, *p < 0.05 vs. native, n = 3–5. **(G)** Group data of relative CH25H mRNA expression in native cells or after 6 h treatment with D39, D39Δ*ply*, D39Δ*cps*, D39Δ*pavA*, or D39Δ*spxB* (each 5×10⁵ CFU/mL), as indicated. Mean ± SEM, *p < 0.05 vs. native, n = 3–5.

Additionally, D39 exposure led to increased expression of ATF3, which is a known transcriptional repressor of CH25H, and downregulation of STAT1, a transcriptional activator of CH25H (Figure 6C-F) ^28,29^. These results suggest that D39 suppresses host antiviral defenses through a coordinated downregulation of CH25H.

To identify the bacterial factors responsible for CH25H suppression, we tested key *S. pneumoniae* virulence mutants, including strains deficient in PLY, capsule, PavA, or SpxB (pyruvate oxidase, producing hydrogen peroxide). However, none of these mutations reversed the downregulation of CH25H or the upregulation of ATF3 (Figure 6G), indicating that these individual factors are not solely responsible.

Because toll-like receptors (TLRs) have been implicated in CH25H induction ^30,31^, we also blocked TLR2 and TLR4 using neutralizing antibodies. Again, neither receptor blockade prevented D39-mediated changes in CH25H, ATF3, or STAT1 expression (Supplementary Figures 2 and 3). LPS-induced TNF-α production was used to confirm TLR blockade efficiency, and these results suggest that neither TLR signaling nor TNF-α are central to the D39-driven CH25H suppression (Supplementary Figures 2 and 3).

It has previously been reported that *S. pneumoniae* binds to and functionally inhibits the PAFR, likely via ChoP moieties present in its teichoic acids ^12,32^. We thus hypothesized that PAFR inactivation by D39 may drive CH25H suppression. Indeed, as PAFR activation is linked to Gq/11-mediated Ca²⁺ release in fura-2–loaded HPMECs stimulation with PAF triggered robust intracellular Ca²⁺ elevation, which was abolished by the PAFR antagonist WEB2086, and blocked by D39 (Figure 7A–D). Importantly, in D39-treated cells Ca²⁺ signals induced by ATP was slightly reduced in comparison to control, however, it still remained, excluding major cytotoxicity as a confounding factor.

**Figure 7.**
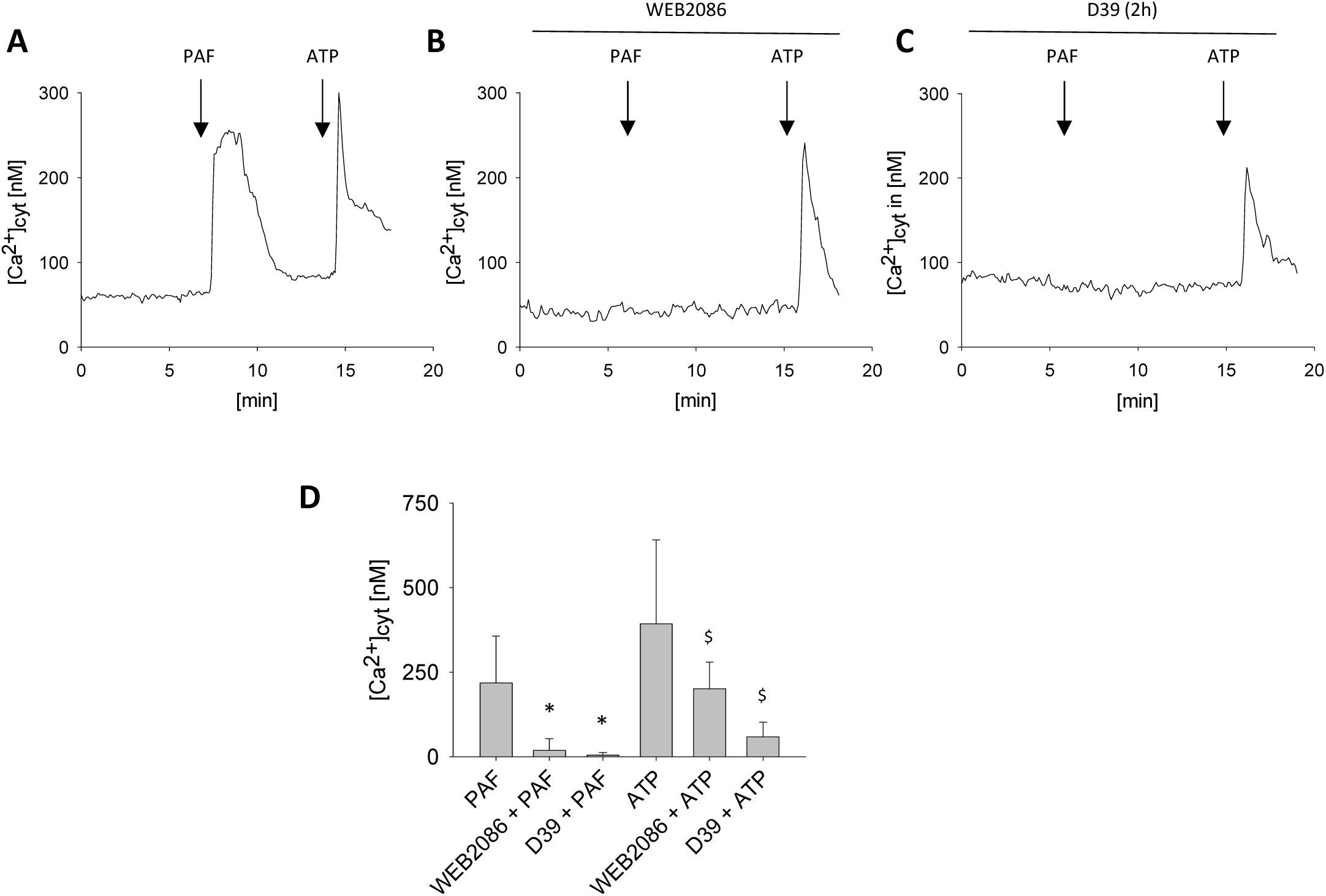
D39 inhibits PAFR. **(A–C)** Tracings of mean endothelial [Ca²⁺]_cyt_ over time under baseline conditions and after application of PAF (10 µM) or ATP (100 µM) in the supernatant of untreated (native) HPMECs or HPMECs pretreated with WEB2086 (10 µM) or D39 (5×10⁵ CFU/mL), as indicated. **(D)** Group data of maximal [Ca²⁺]_cyt_ responses upon PAF (10 µM) or ATP (100 µM) stimulation in native, WEB2086-pretreated, or D39-pretreated HPMECs, as indicated. Mean ± SEM, *p < 0.05 vs. PAF, ^$^p < 0.05 vs. ATP, n = 6–12.

Functionally, both D39 and WEB2086 suppressed CH25H and partially reduced STAT1 expression, with the combination showing additive effects (Figure 8A–B). Notably, neither compound affected the D39-induced upregulation of ATF3 (Figure 8C), suggesting that ATF3 activation is not downstream of PAFR inhibition. Furthermore, neither PAF, WEB2086, nor their combination altered basal ATF3 levels.

**Figure 8.**
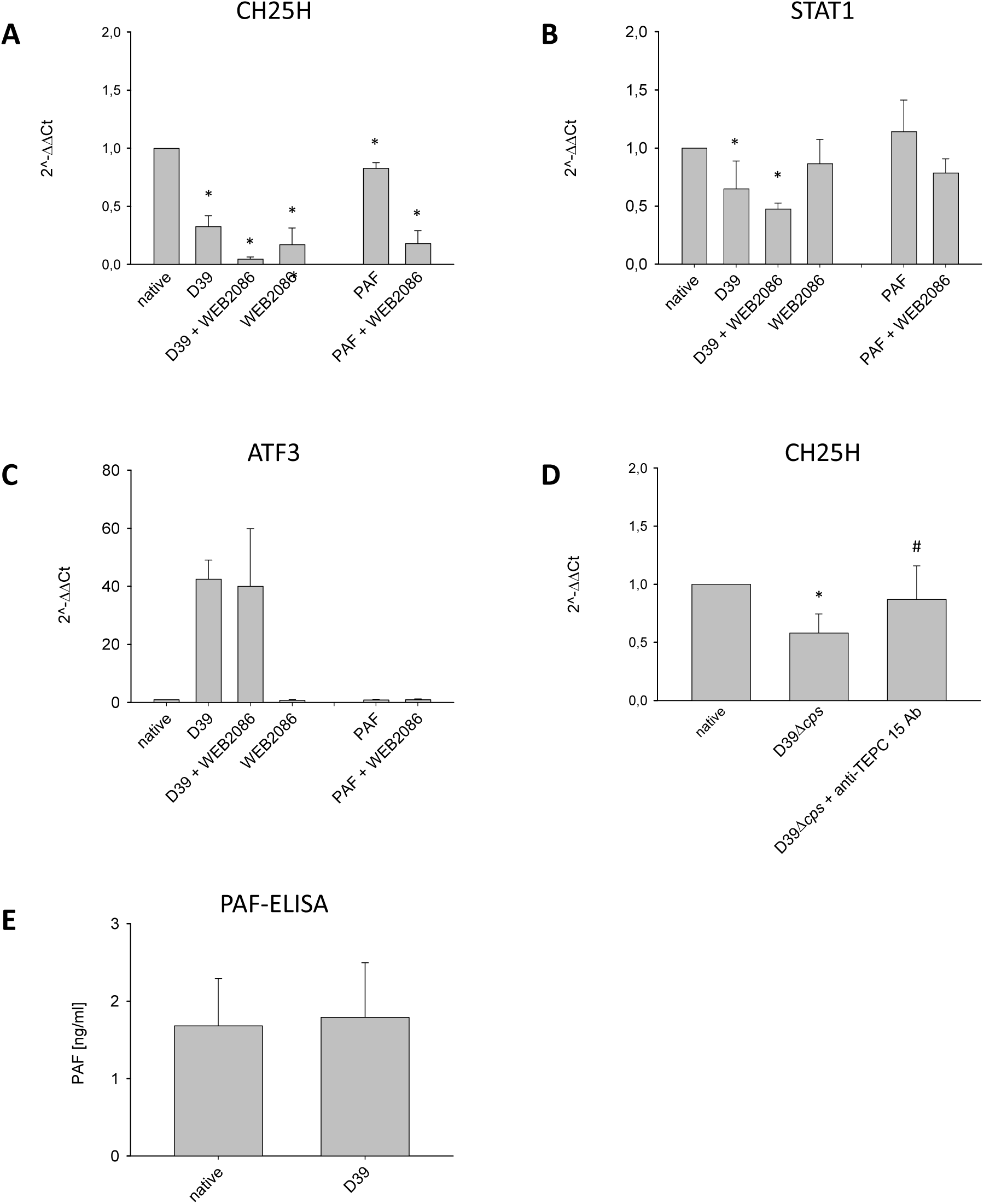
PAFR signaling regulates CH25H expression. **(A–C)** Group data of relative mRNA expression of CH25H, ATF3, and STAT1 in HPMECs. Cells were either untreated (native) or treated with D39 (5×10⁵ CFU/mL), WEB2086 (10 µM), the combination of both, PAF (10 µM) alone, or PAF in combination with WEB2086 for 6 hours, as indicated. Mean ± SEM, *p < 0.05 vs. native, n = 3–4. **(D)** Group data of relative CH25H mRNA expression in HPMECs treated for 6 hours with either D39Δ*cps* (5×10⁵ CFU/mL) alone or in combination with anti-TEPC15 antibody (500 µg/mL), as indicated. Mean ± SEM, *p < 0.05 vs. native, ^#^p < 0.05 vs. D39Δ*cps*, n = 12. **(E)** PAF concentration in the supernatant of HPMECs treated for 6 hours with vehicle or D39 (5×10^5^ CFU/ml), as indicated.

To test whether blockade of bacterial binding to PAFR could rescue CH25H expression, we pretreated a capsule-deficient strain (D39Δ*cps*) with the anti-ChoP antibody TEPC15. This pretreatment abrogated the D39Δ*cps*-induced CH25H downregulation (Figure 8D). We used D39Δ*cps* instead of D39 because the capsule reduces antibody accessibility and binding, and CH25H suppression was similar between both strains (Figure 6G) ^12,33,34^.

Finally, we assessed whether HPMECs produce PAF constitutively, because both D39 and WEB2086 reduced basal CH25H levels. Low-level PAF secretion was confirmed via ELISA (Figure 8E), suggesting a role for endogenous PAFR activation in maintaining CH25H expression.

Taken together, these data indicate that *S. pneumoniae* downregulates CH25H expression in endothelial cells through PAFR inhibition.

### Platelet-activating factor directly antagonizes pneumolysin pore formation independently of PAF receptor signaling

The suppression of CH25H expression was unexpected because 25-HC strongly inhibited PLY-dependent PM pore formation and lung injury in the experiments above. However, inhibition of the CH25H–25-HC pathway could preserve intravascular coagulation and bacterial trapping, suggesting that additional mechanisms may limit toxin activity without blocking vascular responses. Because pneumococci interact with PAFR and PAF signaling is closely linked to platelet activation and inflammatory responses, we asked whether PAF itself influences PLY-induced PM pore formation.

Unexpectedly, we observed that treatment with PAF (10 µM), which was originally applied to test D39-mediated PAFR inhibition (Figure 7A–D), completely abolished D39- and PLY-induced Ca²⁺ oscillations in HPMECs (Figure 9A, B, E, F, I-L). In some cells, PLY stimulation led to an overall elevation of mean [Ca²⁺]_cyt_, consistent with net Ca²⁺ influx exceeding efflux mechanisms until a new steady state was reached (Supplementary Figure 4). Remarkably, even in these cells, PAF application not only eliminated oscillatory Ca²⁺ fluctuations but also normalized the elevated cytosolic Ca²⁺ concentration, highlighting a potent inhibitory effect of PAF on PM pore-mediated Ca²⁺ entry.

**Figure 9.**
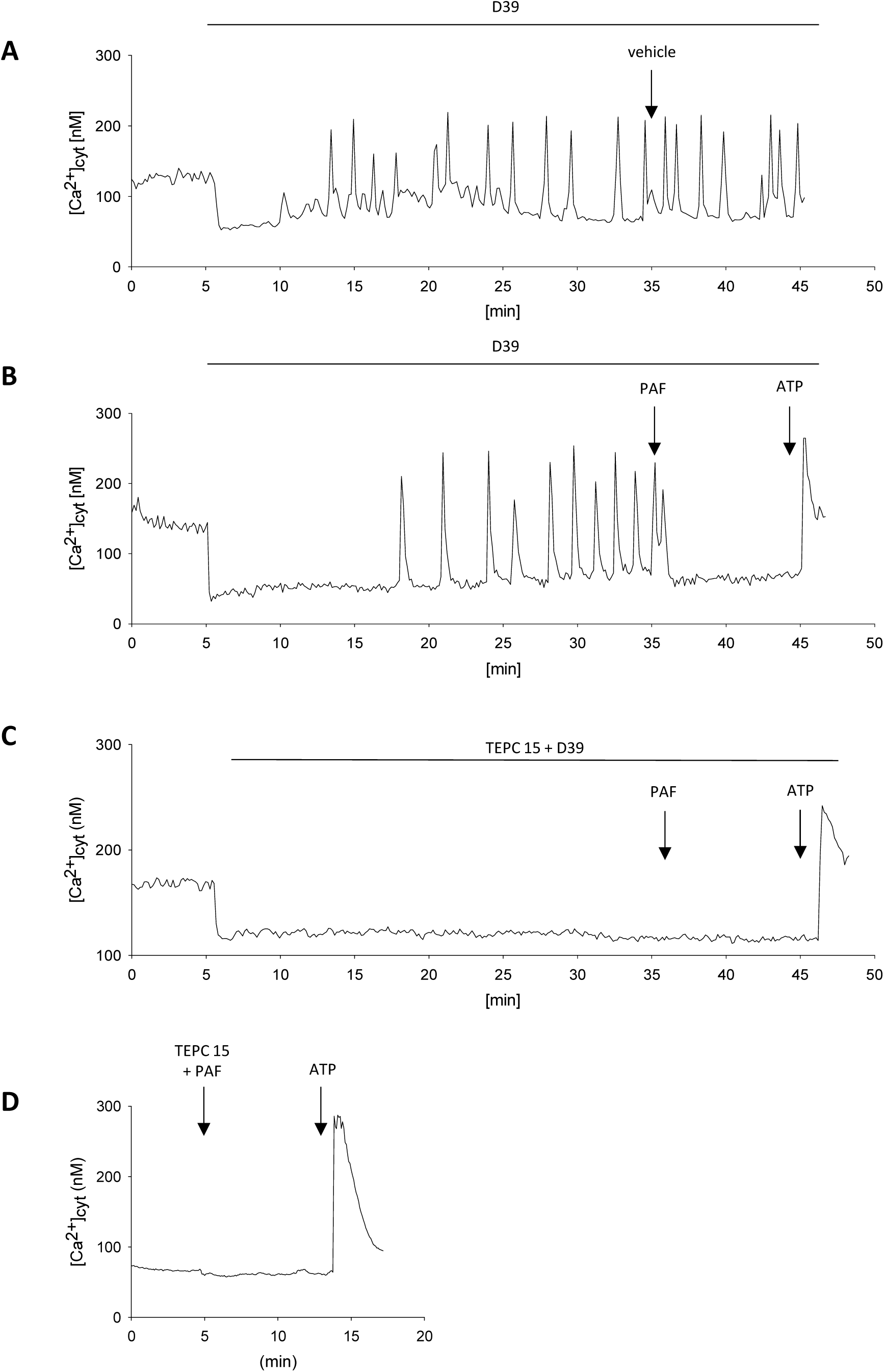
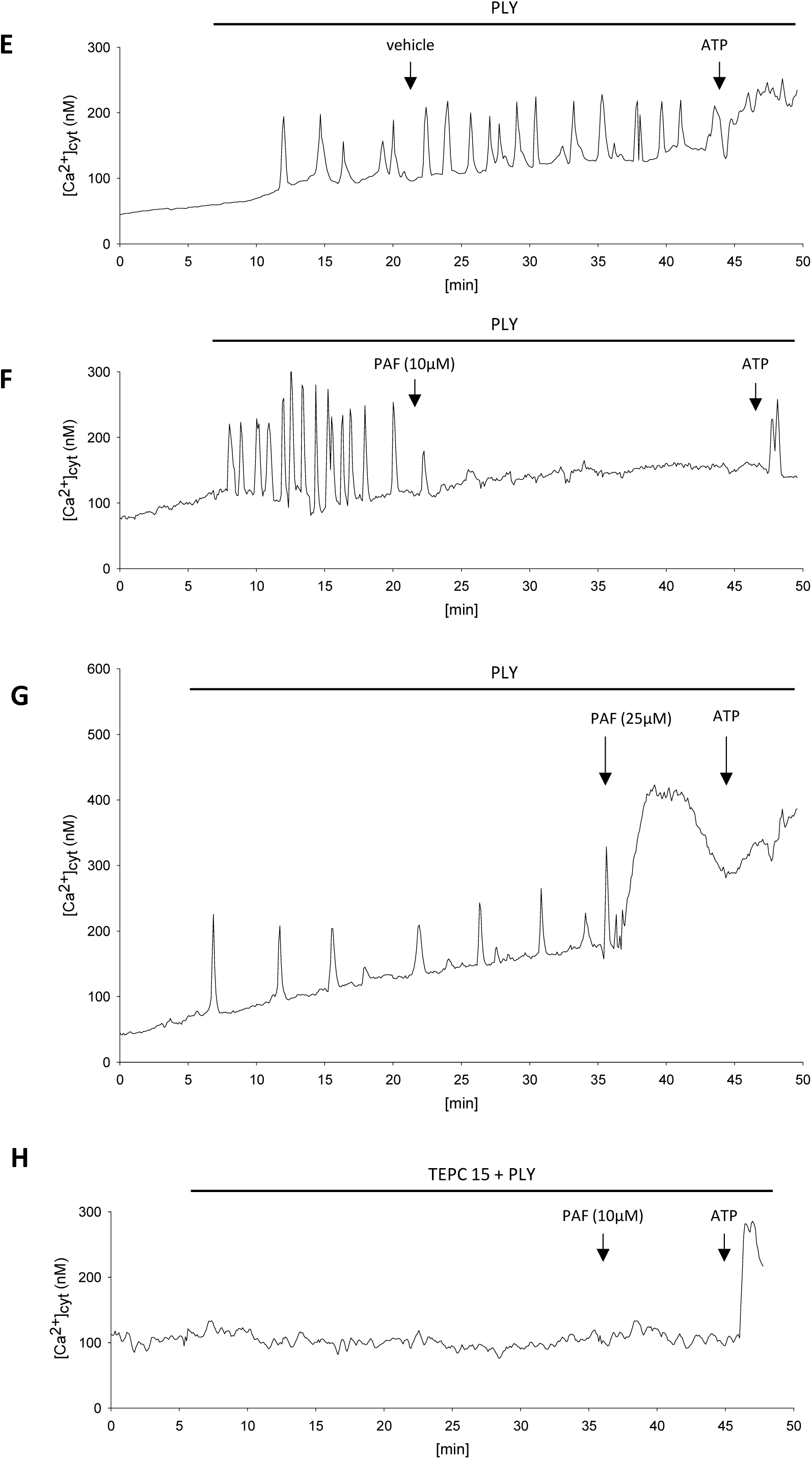
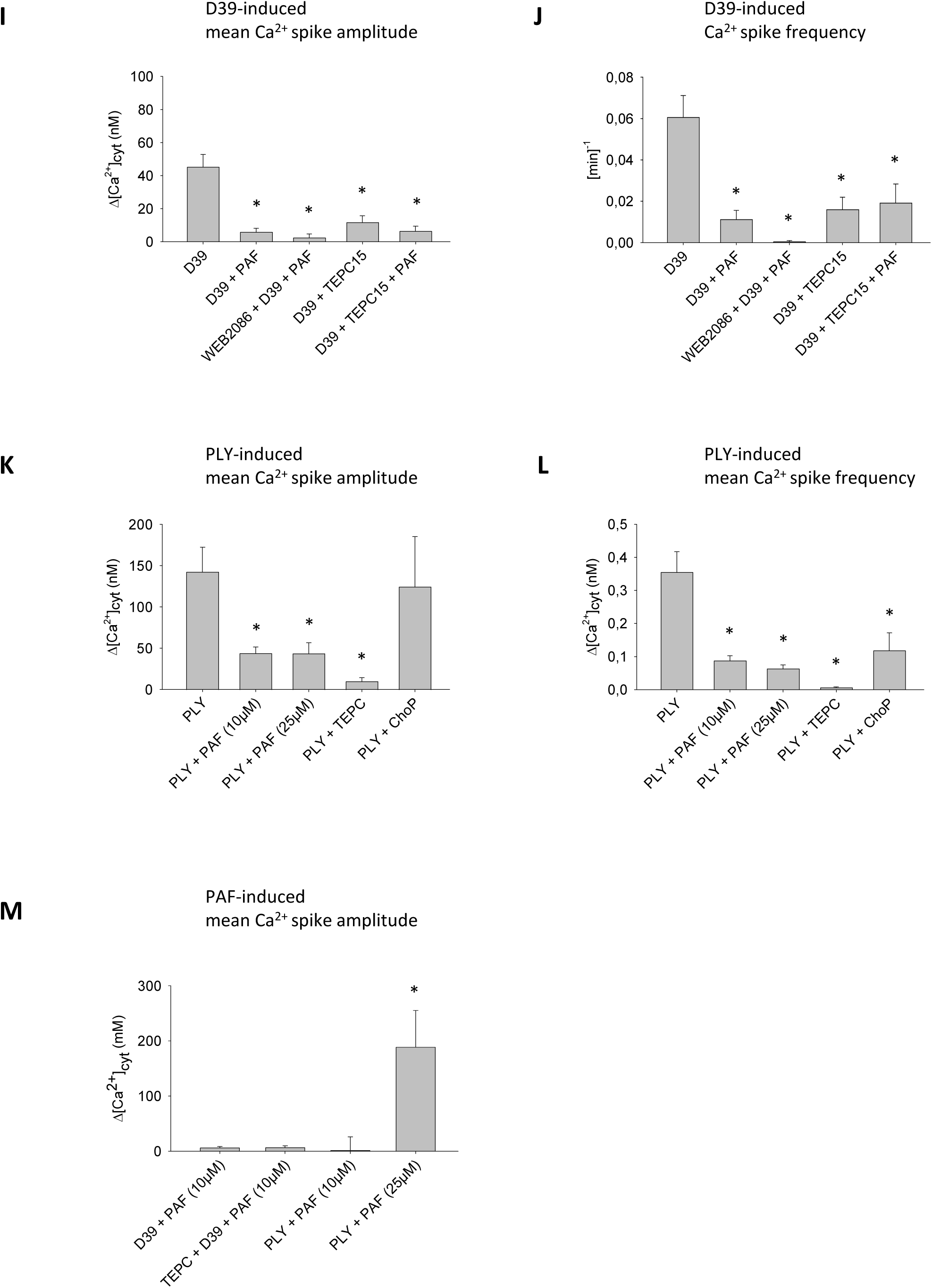
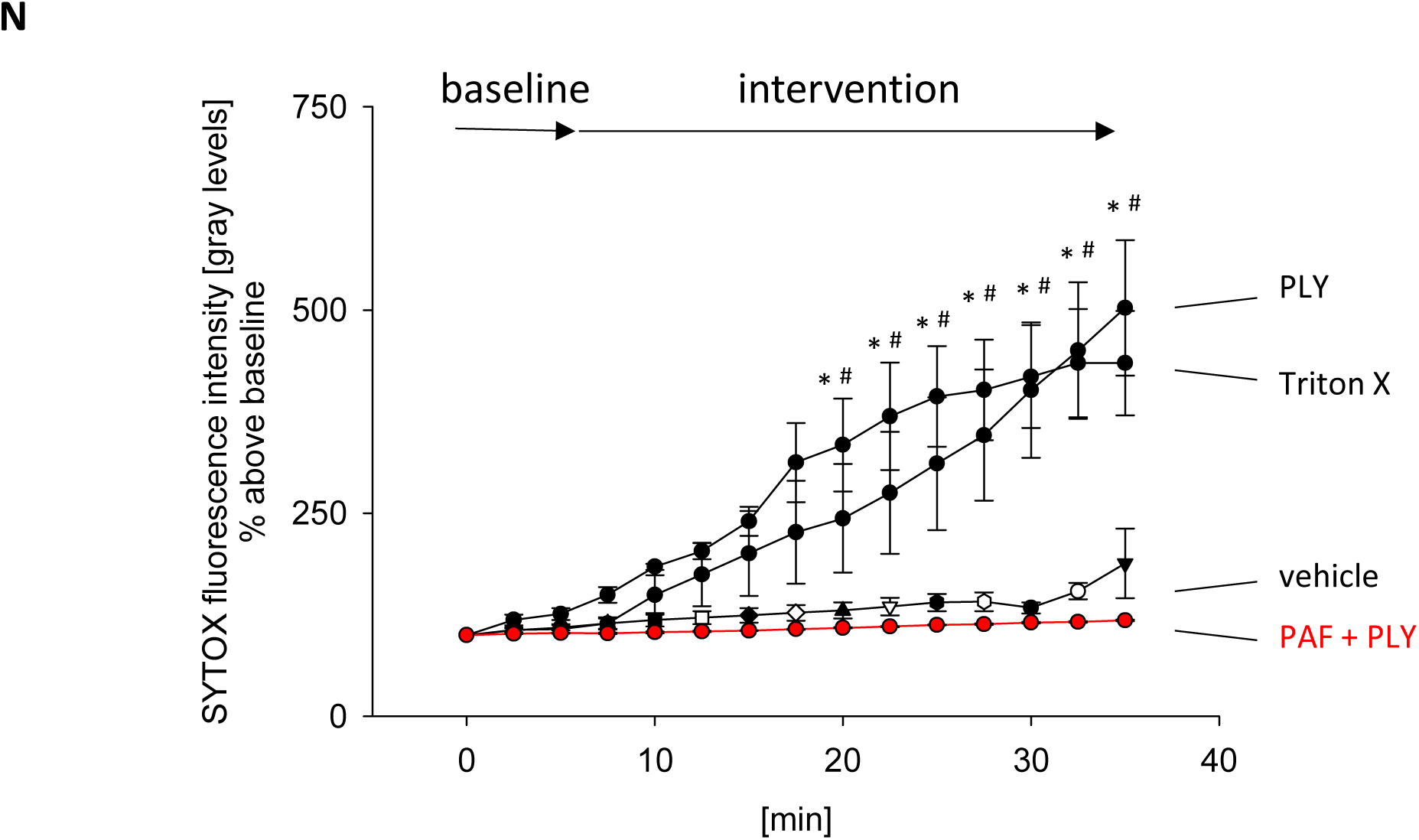
PAF and TEPC15 inhibit PLY-induced Ca²⁺ signaling in HPMECs. **(A–D)** Tracings of endothelial [Ca²⁺]_cyt_ under baseline conditions and following sequential application of D39 (5×10⁵ CFU/mL), D39 (5×10^5^ CFU/mL) pretreated with TEPC15 (500 µg/mL), TEPC15 (500 µg/mL), vehicle, or PAF (10 µM), and ATP (100 µM) in the supernatant of untreated HPMECs, as indicated. **(E–H)** Tracings of endothelial [Ca²⁺]_cyt_ under baseline conditions and following sequential application of PLY (50µg/ml), PLY (50µg/ml) pretreated with TEPC15 (500 µg/mL), vehicle, or PAF (10 µM), and ATP (100 µM) in the supernatant of untreated HPMECs, as indicated. **(I, J)** Group data of mean Ca²⁺ oscillation amplitude (I) and frequency (J) in native HPMECs upon D39 (5×10⁵ CFU/mL) alone or in combination with PAF (10 µM) or WEB2086 (10 µM) or TEPC15 (500 µg/mL), as indicated. Mean ± SEM, *p<0.05 vs. D39, n = 5-11. **(K, L)** Group data of mean Ca²⁺ oscillation amplitude (K) and frequency (L) in native HPMECs upon PLY (50 ng/mL) alone or in combination with PAF (10 or 25 µM), or TEPC15 (500 µg/mL) or phosphocholine (ChoP, 10 µM), as indicated. Mean ± SEM, *p<0.05 vs. PLY, n = 6-12. **(M)** Group data of PAFR-mediated Ca²⁺ response upon PAF (10 or 25 µM), in HPMECs pretreated with D39 (5×10^5^ CFU/mL) alone or in combination with TEPC15 (500 µg/mL), or PLY (50 ng/mL) alone as indicated. Mean ± SEM, *p < 0.05 vs. D39 or PLY, respectively; n = 5–12. Mean ± SEM, *p<0.05 vs. D39 or PLY respectively, n = 5-12. **(N)** Endothelial cells were loaded with SYTOX Green (5 µM) and monitored for fluorescence over time. Baseline was recorded for 5 min before intervention. Exposure to PLY (50 ng/ml) caused a marked increase in SYTOX fluorescence, reflecting plasma membrane permeabilization. A similar increase of SYTOX uptake was obtained by Triton X (0,01 %). This response was completely abolished by co-application of PAF at 10 µM. Data are presented as Mean ± SEM; ^#^p < 0.05 vs. vehicle, *p < 0.05 vs. PAF; n = 3–7.

To corroborate these Ca²⁺ imaging data with an independent readout of PM integrity, we measured SYTOX uptake in endothelial cells. PLY caused a rapid and sustained increase in SYTOX fluorescence, consistent with PM pore formation (Fig. 9N). This effect was fully abolished by PAF at 5 µM, similar to the protection conferred by 25-HC. These results confirm that PAF prevents PLY-induced PM disruption, thereby inhibiting downstream Ca²⁺ signaling.

Importantly, the inhibitory action of PAF on D39- and PLY-induced Ca²⁺ signaling persisted even in the presence of the PAFR antagonist WEB2086 (Figure 9I, J), indicating that this effect is independent of PAF receptor signaling. This observation raised the possibility that PAF might directly interfere with the PM pore-forming activity of PLY rather than acting through a classical G-protein–coupled mechanism.

Structurally, PAF contains a ChoP headgroup similar to phosphatidylcholine. We hypothesized that PAF might inhibit PLY-induced PM pore formation by occupying a ChoP-binding site required for PLY activity. Supporting this, pretreatment of HPMECs with the anti-ChoP antibody TEPC15 completely abolished D39- and PLY-induced Ca²⁺ oscillations (Figure 9C, H, I–L), suggesting that PLY PM pore formation requires not only cholesterol binding but also interaction with PM-associated ChoP.

Interestingly, in HPMECs pretreated with D39 or with D39 plus TEPC15, subsequent PAF application failed to elicit a PAFR-mediated Ca²⁺ response (Figure 9B, C, M). While PAFR blockade by D39 explains the loss of PAF signaling in the first condition, it cannot account for the absence of PAF response when ChoP binding was blocked by TEPC15 (Figure 9C, M). To further dissect this, we repeated the experiment using isolated PLY instead of D39. Under these conditions, PAF also failed to induce a Ca²⁺ signal, indicating that PLY itself can antagonize PAF (Figure 9F, M).

This reciprocal inhibition was further confirmed by dose escalation: increasing the PAF concentration to 25 µM restored the PAFR-mediated Ca²⁺ response in PLY-treated cells (Figure 9G, M). Finally, we could show a direct inhibitory effect of TEPC15 on PAF by demonstrating that in naïve HPMECs, TEPC15 alone did interfere with PAF-induced Ca²⁺ signaling (Figure 9D).

Together, these findings suggest that PAF and PM ChoP compete for PLY. This competition functionally disrupts both PLY-mediated pore formation and PAF-mediated signaling, representing a previously unrecognized antagonistic interaction between a bacterial toxin and a host lipid mediator.

## Discussion

Severe pneumococcal pneumonia and sepsis are frequently associated with acute lung injury, microvascular dysfunction, and high mortality despite appropriate antibiotic therapy. Clinical deterioration often continues after treatment initiation, indicating that toxin-mediated host injury contributes substantially to disease progression ^1,14,15^. In this study, we combined isolated perfused lungs, *in situ* imaging, and cellular experiments to analyze how PLY induces pulmonary vascular injury and how host lipid mediators regulate this response. Our data identify PM pore-dependent Ca²⁺ influx as the central event linking pneumococcal infection to microvascular dysfunction and reveal a lipid-based regulatory system that limits PM damage while preserving antibacterial responses.

Using *in situ* two-photon microscopy, we observed rapid platelet and leukocyte recruitment, perfusion failure, pulmonary edema, and bacterial trapping after perfusion with D39. These responses were largely absent with D39ΔPLY, supporting the concept that PLY is a major driver of vascular injury. The isolated lung preparation allowed direct visualization of microvascular events under controlled conditions, but does not fully reproduce the complexity of systemic infection, and additional host factors may influence toxin effects *in vivo*.

The vascular changes depended on extracellular Ca²⁺ and were accompanied by PM permeabilization and Ca²⁺ influx in endothelial cells and leukocytes *in situ*. Experiments in endothelial cells confirmed that PLY induces SYTOX uptake and Ca²⁺ entry that is abolished in Ca²⁺-free conditions and independent of endogenous Ca²⁺ channels. Mn²⁺ quenching further indicated entry of divalent cations through non-selective PM defects, consistent with pore formation. These findings support the view that toxin-induced PM pores represent the proximal trigger of the inflammatory and hemodynamic responses observed in the lung. However, we cannot exclude that additional pneumococcal factors contribute to vascular dysfunction during infection.

25-HC strongly inhibited PLY-dependent Ca²⁺ influx and protected against vascular leakage and perfusion failure, indicating that PM lipid composition critically determines susceptibility to toxin injury. Because CDCs require accessible PM cholesterol, the protective effect of 25-HC is most consistent with reduced cholesterol availability. In endothelial cells, inhibition of ACAT reversed the effect of 25-HC, supporting a mechanism involving cholesterol esterification ^7–9^. In contrast, protection was also observed in erythrocytes, which lack ACAT, suggesting that oxysterols can additionally alter PM structure directly. Such biophysical effects may redistribute cholesterol within the bilayer and interfere with toxin binding. Although these findings strongly support a lipid-dependent mechanism, the precise structural changes that prevent PM pore formation remain to be defined.

Beyond direct PM injury, PLY triggered complex microvascular responses including platelet adhesion, leukocyte recruitment, perfusion heterogeneity, and bacterial trapping. These processes were Ca²⁺-dependent and largely prevented by 25-HC, indicating that PM pore-mediated Ca²⁺ influx initiates downstream inflammatory and hemostatic reactions. While these responses contribute to vascular dysfunction, they may also represent a host defense mechanism by promoting intravascular sequestration of bacteria. This dual role complicates interpretation, because inhibition of toxin activity may simultaneously protect tissue and impair bacterial containment.

Based on the protective effect of 25-HC, induction of the CH25H–25-HC pathway would be expected during pneumococcal infection. Surprisingly, we observed the opposite: CH25H expression was markedly suppressed. This finding contrasts with viral and interferon-mediated responses, in which CH25H is typically induced ^6,35^, and suggests that pneumococci actively interfere with this pathway. Suppression was independent of several known virulence factors and was reproduced by inhibition of PAF receptor signaling, consistent with the ability of pneumococci to bind PAFR through ChoP-containing teichoic acids ^12,36^. These results indicate that downregulation of CH25H is not a nonspecific stress response but a regulated process.

At first sight, reduced CH25H expression appears disadvantageous for the host, because it increases susceptibility to pore formation. However, suppression of 25-HC may preserve Ca²⁺-dependent vascular activation and coagulation, which contribute to bacterial trapping. This raises the possibility that CH25H downregulation reflects a regulatory trade-off between tissue protection and antibacterial defense rather than simple immune evasion.

The suppression of CH25H raised the question of how PLY activity is controlled when endogenous 25-HC production is reduced. Because inhibition of the CH25H–25-HC pathway may preserve coagulation-dependent bacterial trapping but simultaneously increases susceptibility to PM damage, additional mechanisms must exist that limit PM pore formation without abolishing vascular activation. PAF was of particular interest in this context, because pneumococci interact with the PAFR through ChoP-containing cell wall components, and PAF signaling is closely linked to platelet activation and inflammatory responses ^13,37,38^.

Unexpectedly, we found that PAF itself inhibited PLY-induced Ca²⁺ influx and PM permeabilization. This effect persisted in the presence of a PAFR antagonist, indicating that PAF does not act through classical receptor signaling but interferes directly with toxin activity. The observation that the ChoP-specific antibody TEPC15 produced a similar inhibition, and that PLY and PAF mutually interfered with each other’s signaling, suggests that PM pore formation requires interaction not only with cholesterol but also with ChoP-containing PM components. PAF may therefore act by competing for this interaction site, thereby preventing efficient toxin binding or oligomerization. Although the precise molecular mechanism remains to be clarified, these findings identify a previously unrecognized lipid-dependent step in PLY PM pore formation.

The ability of PAF to inhibit pore formation provides a possible explanation for the paradoxical downregulation of CH25H during pneumococcal infection. Reduced 25-HC levels allow Ca²⁺-dependent vascular activation and intravascular aggregation, which may facilitate bacterial trapping, whereas PAF could limit excessive PM damage without suppressing these responses. In this model, distinct lipid mediators regulate different aspects of the host response: 25-HC primarily protects PM integrity by reducing accessible PM cholesterol, whereas PAF interferes with ChoP-dependent toxin binding while preserving platelet activation and coagulation. This dual regulation may allow the host to balance tissue protection with antibacterial defense under conditions of high toxin exposure.

The interaction between pneumococci and the PAFR adds an additional layer of complexity. Binding of bacterial ChoP to PAFR inhibits receptor signaling and contributes to CH25H suppression, but at the same time circulating PAF may still antagonize PLY independently of the receptor. When PAFR signaling is restored, PAF can again promote platelet activation and endothelial responses while simultaneously limiting pore formation. Such context-dependent separation of receptor-dependent and receptor-independent effects may represent a flexible mechanism that allows the host to adapt to different stages of infection.

These findings may have implications for the pathogenesis and treatment of pneumococcal sepsis. Antibiotic therapy causes bacterial lysis and can lead to massive release of PLY, potentially aggravating tissue injury even while bacterial load decreases ^39,40^. Strategies aimed at neutralizing PLY could therefore reduce organ damage, but complete inhibition of PM pore formation might impair intravascular trapping of bacteria. Modulation of PM lipid composition, for example by oxysterols, may protect the endothelium but could also reduce coagulation-dependent containment. In contrast, interference with ChoP-dependent toxin binding might limit PM pore formation without fully suppressing vascular activation. Whether such approaches are beneficial *in vivo* remains uncertain and will require further investigation.

Although the present study focused on PLY, the regulatory principles identified here may extend to other CDCs and related PM pore-forming toxins ^9,41^. Many Gram-positive pathogens produce toxins that interact with PM cholesterol, and the availability of accessible cholesterol and ChoP-containing lipids may be critical determinants of toxin susceptibility. Host regulation of PM lipid composition could therefore represent a general mechanism for controlling PM pore-forming toxins while preserving immune function.

In summary, our data identify PLY-induced PM pore formation as the central trigger of pulmonary vascular injury and demonstrate that host lipid mediators regulate toxin activity in a bidirectional manner. Suppression of the CH25H–25-HC pathway permits Ca²⁺-dependent vascular activation and bacterial trapping, whereas PAF provides a complementary mechanism that limits excessive PM damage. This lipid-based control of PM susceptibility offers a framework for understanding how the host balances tissue protection with antibacterial defense during pneumococcal infection and may suggest new approaches to limit toxin-mediated injury without compromising host immunity.

## Materials and Methods

### Materials

Fluorophores Fura-2 AM and Fluo-4 AM were dissolved in 0.02% Pluronic F-127 (both from PromoKine, Heidelberg, Germany). SYTOX Green (dissolved in dimethyl sulfoxide) was obtained from Thermo Fisher Scientific (Rockford, IL, USA). ChoP was purchased from MedChemExpress (Monmouth Junction, NJ, USA). Sandoz 58-035, Mn^2+^ chloride tetrahydrate, gadolinium (III) chloride hexahydrate, 25-HC, β-Acetyl-γ-O-hexadecyl-L-α-phosphatidylcholine hydrate (PAF), murine IgA kappa monoclonal antibody (clone TEPC15), and thapsigargin were acquired from Sigma-Aldrich Chemie GmbH (Taufkirchen, Germany). SMase from *Staphylococcus aureus* and WEB-2086 were purchased from Enzo Life Sciences GmbH (Lörrach, Germany). 7α-25-HC was from Cayman Chemicals (Ann Harbor, Mi, USA). GW 3965 hydrochloride, T 0901317, and GSK 2033 were purchased from Tocris Bioscience (Wiesbaden, Germany). Unless otherwise indicated, all solutions and reagents were prepared in Hank’s Balanced Salt Solution (HBSS; 150 mM Na⁺, 5 mM K⁺, 1 mM Ca²⁺, 1 mM Mg²⁺, and 20 mM HEPES; pH 7.4). For experiments under Ca²⁺-free conditions, Ca²⁺-free HBSS was used.

### Cell culture

HPMECs were purchased from PromoCell GmbH and cultured in Endothelial Cell Growth Medium MV (PromoCell GmbH, Heidelberg, Germany) at 37 °C in a humidified atmosphere with 5% CO₂.

### Bacteria and culture

The following *Streptococcus pneumoniae* strains were used: wild-type D39 (D39, serotype 2; NCTC 7466), capsule-deficient D39Δ*cps*, PLY-deficient, D39Δ*ply*, PavA-deficient D39Δ*pavA*, and pyruvat-deficient, D39Δ*spxB*. All strains have been published earlier ^42–45^. Bacteria were plated on blood agar and grown at 37 °C for 16 hours. Colonies were then resuspended in THY medium (Todd Hewitt broth supplemented with 5% yeast extract) and cultured at 37 °C for 2–16 hours. After growth, bacterial cultures were harvested and stored in THY medium containing 20% glycerol at −80 °C at a concentration of 1 × 10⁷ CFU/mL. For selection of mutant strains, kanamycin (30 µg) was used for D39Δ*cps*, erythromycin (15 µg) was used for the other mutants, and, applied via antibiotic-impregnated paper discs (Becton Dickinson, Sparks, MD, USA). For experiments involving PLY, recombinant, cytolytically active PLY was used, produced as previously described ^46^.

### Real-Time Quantitative RT-PCR (RT-qPCR)

Total RNA was isolated using the RNeasy Kit (Qiagen, Hilden, Germany). For cDNA synthesis, 2 µg of RNA were reverse-transcribed using the Omniscript Reverse Transcription Kit (Qiagen) according to the manufacturer’s instructions. RT-qPCR was performed using gene-specific primers for the following targets:

• Cholesterol-25-hydroxylase

Forward: ACATCTGGCTTTCCGTGGAG

Reverse: TACGGAGCGAAGTTGCAGTT

• ATF3

Forward: AGGTCTCTGCCTCGGAAGT

Reverse: CTTCTTCAGGGGCTACCTCG

· STAT1

Forward: CACAAGGTGGCAGGATGTCT

Reverse: GGTGAACCTGCTCCAGGAAT

For normalization, the housekeeping gene GPR138 was used:

Forward: TGGGCTCGTGGGAAACTCAC

Reverse: GTGTAGGCAAAGCGGTGTTA

Primers were designed to span exon–exon junctions to prevent amplification of genomic DNA, except for CH25H, which is encoded by an intronless gene. qPCR amplification was performed on a Rotor-Gene 3000 system (Corbett Research, Mortlake, Victoria, Australia) according to the manufacturer’s instructions. Relative gene expression was calculated using the ΔΔCt method.

### Western Blotting

Cells were lysed in RIPA buffer supplemented with a protease inhibitor cocktail (Complete Mini, EDTA-free; Roche, Mannheim, Germany) for 30 minutes on ice. Lysates were centrifuged at 12,000 × g for 20 minutes at 4 °C. Whole-cell lysates (20 µg) were mixed with sample buffer (NuPAGE Sample Buffer, Life Technologies GmbH, Darmstadt, Germany) and reducing agent (NuPAGE Sample Reducing Agent, Life Technologies GmbH), then heated at 95 °C for 5 minutes. Proteins were separated by SDS-PAGE on a 4–12% Bis-Tris polyacrylamide gel (NuPAGE, Life Technologies GmbH) and transferred to nitrocellulose membranes via electroblotting.

Membranes were blocked in PBS containing 0.1% Tween-20 and 5% milk powder for 1 hour at room temperature with agitation. Subsequently, membranes were incubated overnight at 4 °C with primary antibodies against CH25H (1:200), ATF3 (1:500), and STAT1 (1:500) (all from Thermo Fisher Scientific). After washing, bound antibodies were detected using horseradish peroxidase (HRP)-conjugated secondary antibodies (incubated for 1 hour at room temperature with agitation) and enhanced chemiluminescent substrate (ECL Plus Western Blotting Detection Reagents, GE Healthcare, Freiburg, Germany), according to the manufacturer’s instructions.

For molecular weight reference, the MagicMark™ XP protein ladder (Life Technologies) was used. ß-actin (1:10,000; Merck KGaA, Darmstadt, Germany) served as the loading control.

### Protein Determination

Protein concentrations were measured using the bicinchoninic acid (BCA) protein assay kit (Thermo Fisher Scientific) according to the manufacturer’s instructions.

### Hemolysis Test

Ten milliliters of fresh human blood (collected 1:10 in sodium citrate buffer) were centrifuged at 1,000 × g for 20 minutes at room temperature. The resulting erythrocyte pellet was resuspended three times in 20 mL PBS, each followed by centrifugation at 1,000 × g for 20 minutes. The final erythrocyte suspension was diluted 1:50 in PBS.

Standard hemolysis reaction mixtures (200 µL total volume) were prepared in 500 µL tubes and contained:

- 100 µL lysis buffer (with or without 25-HC at 50 µM),
- 50 µL recombinant PLY (50 ng/mL), PBS (negative control) or Triton X (0.2%, positive control)
- 10 min incubation and then adding
- 50 µL of the 1:50 erythrocyte suspension.

Samples were incubated for 30 minutes at 37 °C, then centrifuged at 1,000 × g for 10 minutes. A total of 100 µL of supernatant from each sample was transferred to a clear, flat-bottom 96-well plate (Thermo Fisher Scientific). Hemolysis was quantified by measuring the absorbance of released hemoglobin at 540 nm using a microplate reader (Thermo Fisher Scientific).

### In vitro live cell imaging

#### Conventional fluorescent microscopy

For fluorescent imaging, a coverslip with fluorophore-loaded HPMEC was mounted in an imaging chamber on the stage of a conventional epifluorescence microscope (Olympus, Hamburg, Germany). Illumination from a mercury lamp directed through appropriate interference filters (Semrock, Rochester, NY, USA) excited the fluorophores. The exposures of fluorophores were controlled by a filter wheel (Sutter Lambda 10-C, Sutter Instrument Co., Novato, CA, USA). The fluorescence emission was collected using an objective lens (40x water immersion, numerical aperture 0.8, Zeiss, Germany) and captured with a charge-coupled device camera (Photometrics Coolsnap HQ2).

#### [Ca2+]_cyt_ determination

For [Ca^2+^]_cyt_ measurements, cells were loaded for 40 min with Fura-2-AM (10 μM) in physiological solution containing 0.02% pluronic. The cells were excited at 340, 380, or 360 nm. The fluorescence emissions at 510nm were recorded and [Ca^2+^]_cyt_ was calculated from a computer-generated 340:380 emissions ratio based on a dissociation constant of 224nmol/l and appropriate calibration parameters ^47^.

#### Pore formation determination

SYTOX green staining: To determinate pore-formation, 5 µM SYTOX green was added to the supernatant of HPMECs and increase of fluorescence (excitation: 488 nm, emission: 523 nm) was recorded continuously over 30 min. SYTOX green is impermeant to live cells with an intact plasma membrane but permeable to leaky membranes and binds nucleic acids. At the end of each experiment 0.01% Triton X was added to the cells as a positive control.

Manganese II (Mn^2+^) Quenching: To evaluate pore formation and thus external Ca^2+^ entry Mn^2+^ quenching (0.5 mM) was conducted in Ca^2+^-containing and Ca^2+^-free buffer of fura-2 loaded cells. Fura-2 emission fluorescence was recorded at excitation wavelength of 360 nm, the isobestic point, where the fura-2 fluorescence absorbance (at 510 nm) is independent from calcium concentration.

### Isolated Blood-Perfused Mouse Lung Model

#### Animals

C57BL/6J (C57BL) mice were purchased from The Jackson Laboratory (Bar Harbor, ME, USA). All animal procedures were reviewed and approved by the local ethics committee and the governmental authorities of Hamburg, Germany. Experiments were conducted in accordance with the “Principles of Laboratory Animal Care” (NIH Publication No. 86-23, revised 1985) and German legislation on animal protection.

#### Lung Preparation

Experimental details have been described previously ^48^. In brief, animals were anesthetized with 5 vol% isoflurane followed by intraperitoneal injection of ketamine (120 mg/kg body weight) and xylazine (25 mg/kg body weight; Rompun™). After tracheotomy and tracheal cannulation, mice were ventilated under volume-controlled conditions using a rodent ventilator (Model 683, Harvard Apparatus, Holliston, MA, USA). The abdomen was opened, and exsanguination was performed via right heart puncture. Cannulas were inserted into the main pulmonary artery (via the right ventricle) and into the left atrium. The heart–lung block was removed *en bloc* and placed on a Plexiglas dish under buffer to prevent drying.

The pulmonary vasculature was perfused using a peristaltic pump with HEPES-buffered solution containing 5% fetal bovine serum at 37 °C. Pulmonary arterial and left atrial pressures were maintained at 10 and 2 cm H₂O, respectively, with a flow rate of 1 mL/min. Lungs were continuously inflated with a gas mixture of 30% O₂, 5% CO₂, and 65% N₂ at an airway pressure of 5 cm H₂O.

### In Situ Live Cell Imaging

#### Setup and Positioning

Isolated, buffer-perfused mouse lungs were placed upside-down to allow microscopy from the basal lung surface.

#### Leukocyte and Platelet Infusion

For leukocyte analysis, 5 mL of whole blood was centrifuged, and the platelet-rich plasma was separated. The buffy coat was incubated with Rhodamine 6G (0.05% solution; 15 µL per 1 mL blood; Thermo Fisher Scientific), centrifuged (2,000 × g, 10 min), and resuspended in Ca²⁺-free HEPES buffer at a final concentration of 0.5×10⁹ leukocytes/mL. The suspension was reinfused just prior to imaging ^48^.

PLTs were isolated from 5 mL of whole blood mixed with 1.5 mL phosphate-buffered saline (PBS) containing 15.2 µM citric acid, 30 µM trisodium citrate, 40 µM dextrose, and 1.5 µg/mL prostaglandin E1 (PGE1). Following centrifugation, 500 µL of platelet-rich plasma was collected and centrifuged again (2,000 × g, 10 min). The resulting pellet was incubated with Calcein red orange (10 µM) for 15 minutes at room temperature. PLTs were washed and resuspended in 2 mL HEPES buffer (0.2 × 10⁹ PLTs/mL) and reinfused prior to imaging ^48^

To visualize resident leukocytes, anti-LY6G-PE antibody (2 µg/mL; Thermo Fisher Scientific) was added to the perfusate 15 minutes before the start of imaging.

For visualization of microhemodynamics, native red blood cells (RBCs) were added to the perfusate, and FITC-albumin was co-perfused to create negative contrast ^48^

#### Conventional Fluorescence Microscopy

Fluorescent imaging was performed using an epifluorescence microscope (Axiotech, Zeiss, Göttingen, Germany) with excitation from a mercury arc lamp filtered through appropriate interference filters (Semrock, Rochester, NY, USA). Fluorophore exposure was controlled via a filter wheel (Sutter Lambda 10-C, Sutter Instrument Co., USA). Emitted fluorescence was collected using a 40× water-immersion objective (NA 0.8; Zeiss, Germany) and recorded with a CCD camera (Photometrics Coolsnap HQ2, USA). Images were captured every 10 seconds and analyzed using MetaFluor software (Molecular Devices, Downington, PA, USA).

#### [Ca²⁺]_cyt_ Measurements

Endothelial [Ca²⁺]_cyt_ levels were measured in subpleural arterioles using the Fura-2 AM method, as previously described ^48,49^. Fura-2 AM (10 µM) was vascularly perfused for 30 minutes. Excitation was performed at 340 nm and 380 nm, and emissions were recorded at 510 nm. [Ca²⁺]_cyt_ were calculated from the 340:380 ratio using a dissociation constant of 224 nM and standard calibration parameters ^47^.

#### Two-Photon Confocal Microscopy

Two-photon imaging was performed on endothelial cells of subpleural pulmonary arteries using a single-beam two-photon microscope as previously described ^50^. The Ti:Sa pulsed laser (MaiTai, Spectra-Physics) was tuned to 750 nm and directed through a 20× water-dipping lens (NA 0.95; working distance 2 mm; Olympus, Hamburg, Germany). Emission photons passed through a 475/47 nm band-pass filter and were detected using photomultiplier tubes (LaVision Biotec, Bielefeld, Germany).

For Ca²⁺ imaging, endothelial cells and resident leukocytes were loaded with Fluo-4 (10 µM) 30 minutes before the experiment. Fluorescence intensity was measured and analyzed.

#### Quantification of Microvascular Parameters

RBC velocity was determined by measuring the mean velocity of 30 FITC-labeled red blood cells traversing a defined vascular region using ImageJ software (v1.53t, NIH).

Leukocyte and platelet kinetics were assessed by defining rolling cells as those traveling at less than 50% of mid-vessel velocity. Adherent cells were defined as those stationary for ≥30 seconds.

Endothelial barrier function was evaluated by measuring FITC-albumin fluorescence within vessels and in surrounding interstitial or alveolar compartments.

### Lung wet/dry ratio

To assess pulmonary edema formation, we determined lung tissue wet/dry weight ratio by drying samples at 100°C for 24 h.

### PAF ELISA

The PAF concentration in the supernatant of HPMECs native or pretreated with D39 (5×10^5^ CFU/ml) was determined with the PAF ELISA KIT (Avia Systems Biology, San Diego, CA, USA) according to the manufacturer’s protocol.

### Statistical Analysis

All data are presented as means ± standard error (SEM). Statistical analysis was performed using SigmaStat software (Systat Software, San Jose, CA, USA). Group comparisons were evaluated using analysis of variance on ranks followed by pairwise multiple comparisons. For repeated measurements, the Mann–Whitney rank-sum test was applied. A p-value < 0.05 was considered statistically significant.

## Acknowledgement

We wish to thank Kirsten Pfeiffer-Drenkhahn, Andrea Pawelczyk, Claudia Luechau, and Monika Weber for their great technical assistance. Without it, this work would not have been possible.

## Funding

This study was funded by grants of the Deutsche Forschungsgemeinschaft to R.K. (KI 867/7-1) and S.H. (HA 3125/8-1; project number 523973396) and Johanna und Fritz Buch-Gedächtnis-Stiftung.

## Author contributions

Conceptualization, methodology, investigations and writing of manuscript by R. K. and M. K.; Resources, review and editing of the manuscript by S. H. and A. D. Funding acquisition by R. K.

## Competing interests

The authors declare no competing interests.

---

**Supplementary figure 1:**
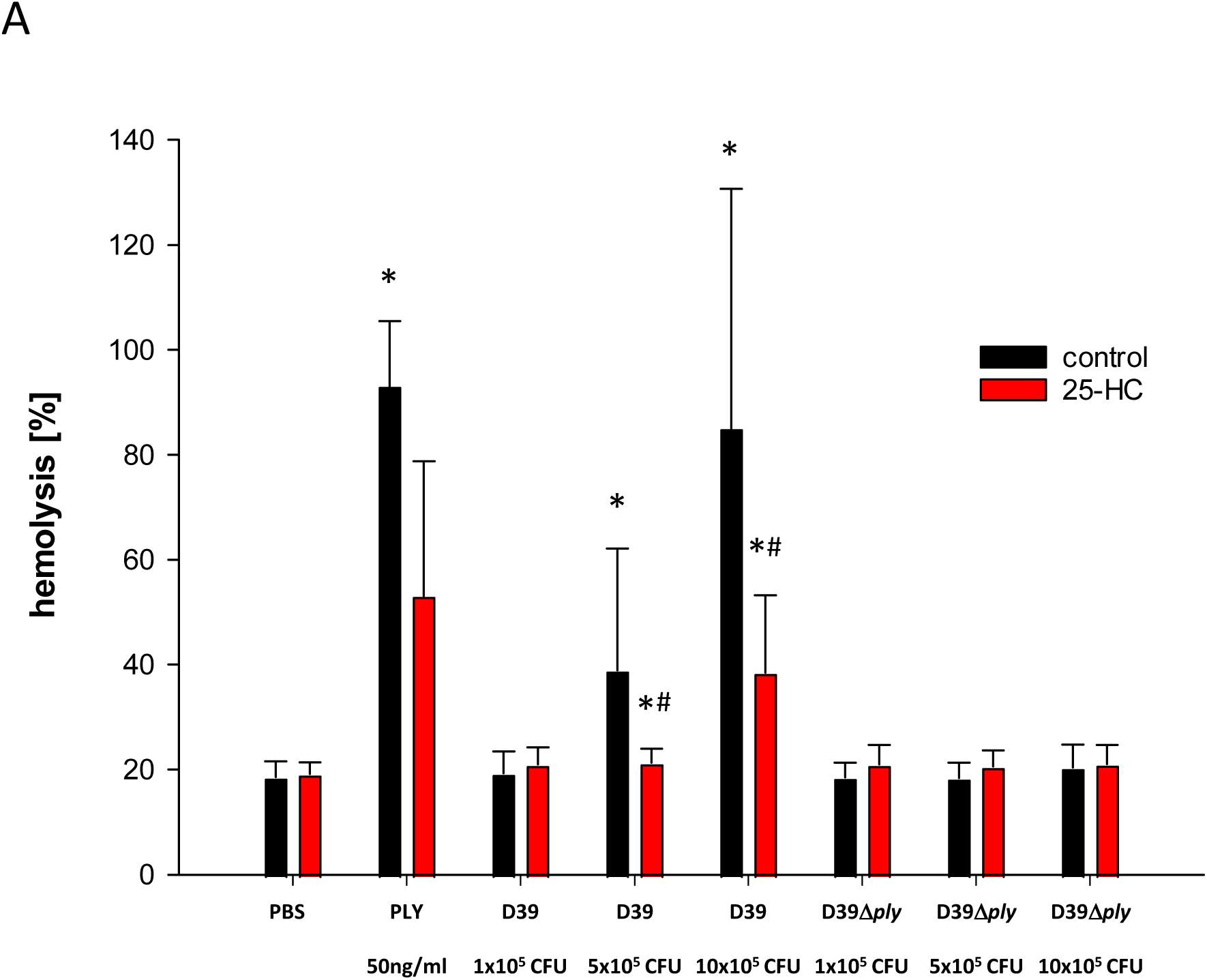
25-HC inhibits PLY-mediated red blood cell lysis and plasma membrane pore formation in HPMECs. Quantification of hemolysis induced by PLY (50 ng/mL), D39 (10^5^, 5×10^5^, or 10×10^5^ CFU/mL), or the PLY-deficient mutant D39Δ*ply* (10^5^, 5×10^5^, or 10×10^5^ CFU/mL) in human red blood cells (RBCs). Hemolysis is expressed as a percentage of maximal lysis. Pretreatment with 25-HC (50 µM) abolished PLY- and D39-induced hemolysis. Data are presented as Mean ± SEM; *p < 0.05 vs. PBS, ^#^p < 0.05 vs. PLY or D39 without 25-HC; n = 8.

**Supplementary Figure 2.**
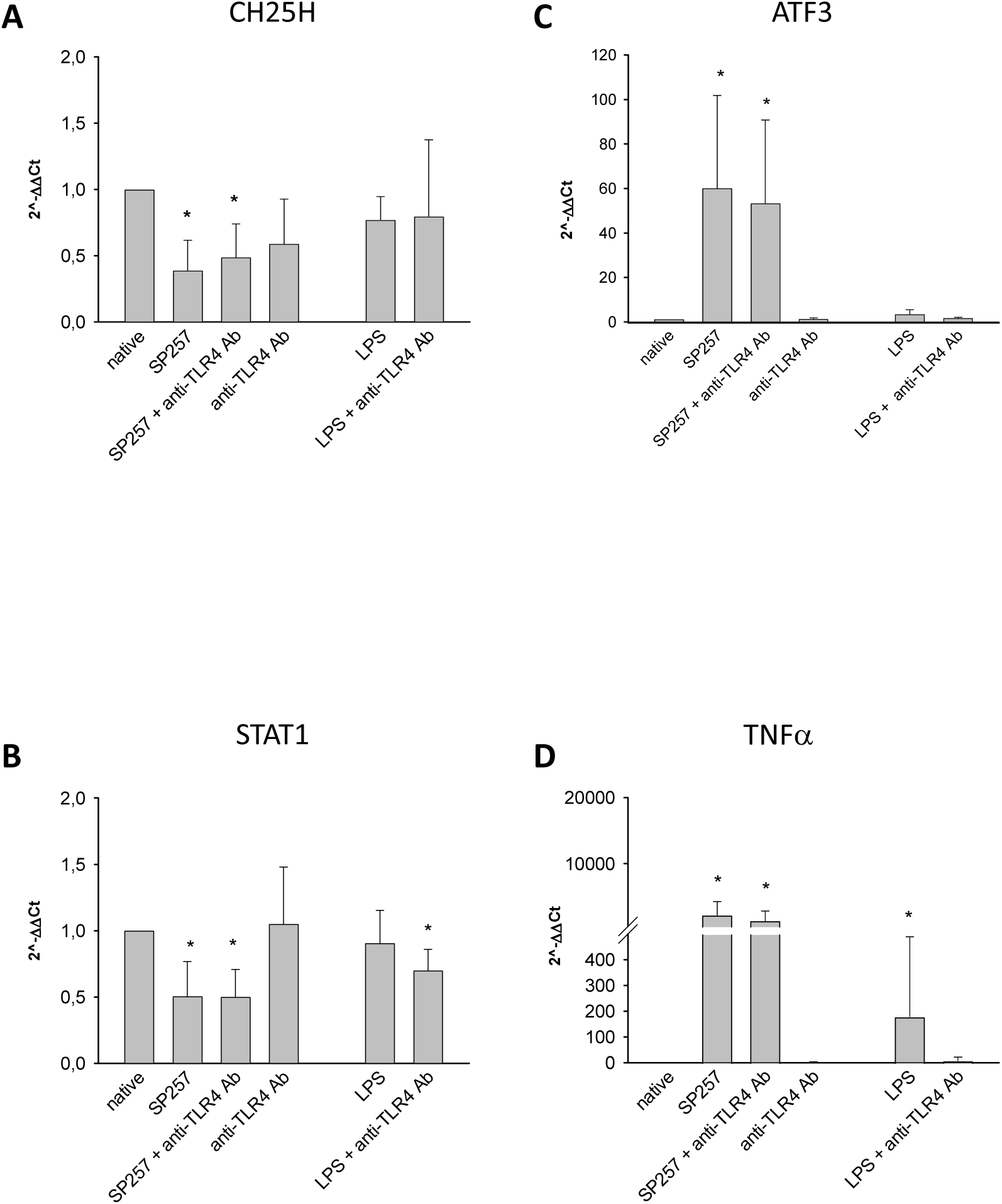
TLR4 does not regulate CH25H expression. **(A–D)** Group data of relative mRNA expression of CH25H, ATF3, STAT1, and TNFα in HPMECs. Cells were either untreated (native) or treated with D39 (5×10⁵ CFU/mL), anti-TL4 Ab (10 µM), the combination of both, LPS (10 µM) alone, or LPS in combination with anti-TL4 Ab for 6 hours, as indicated. Mean ± SEM, *p < 0.05 vs. native, n = 3–4.

**Supplementary Figure 3.**
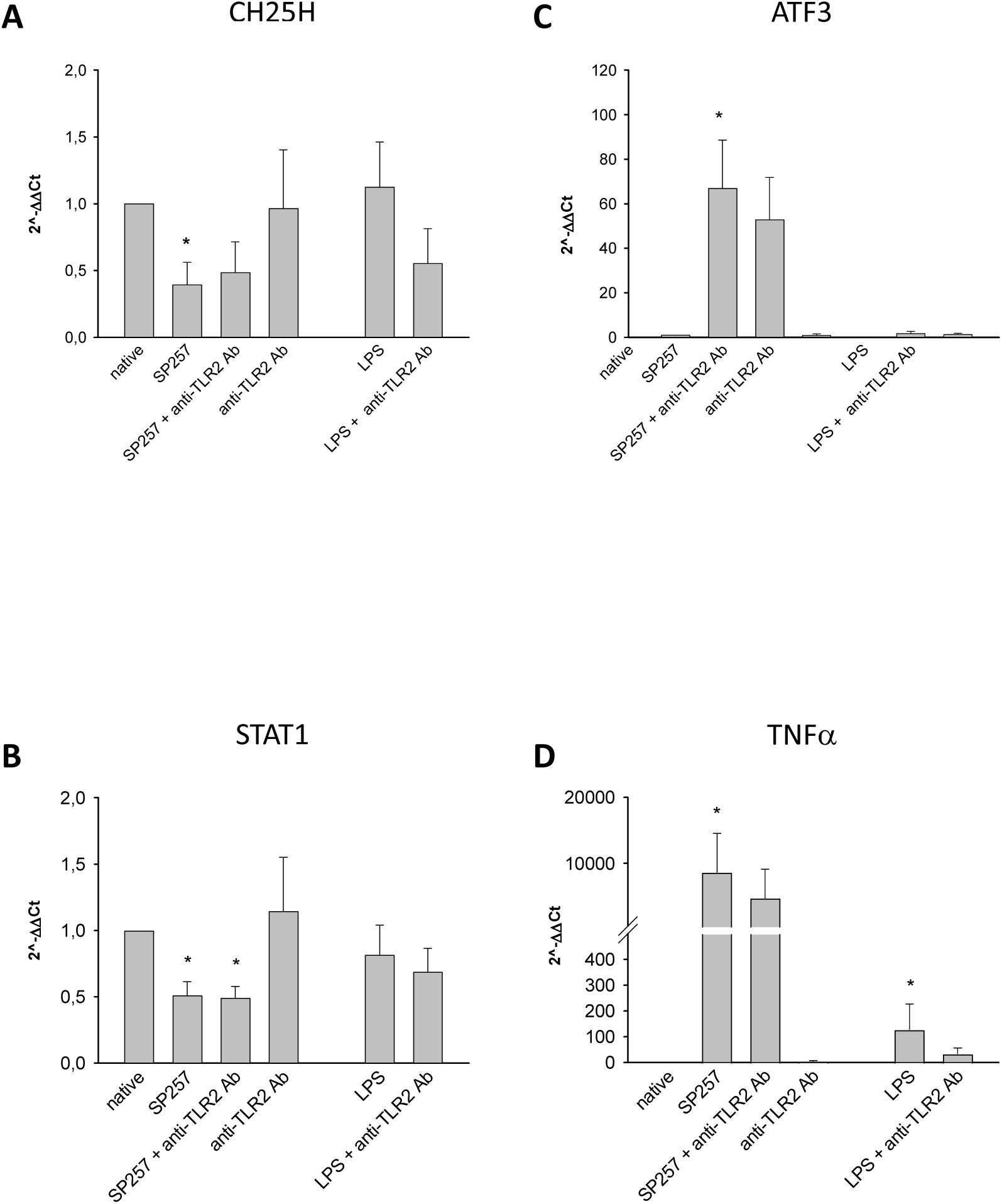
TLR2 does not regulate CH25H expression. **(A–D)** Group data of relative mRNA expression of CH25H, ATF3, STAT1, and TNFα in HPMECs. Cells were either untreated (native) or treated with D39 (5×10⁵ CFU/mL), anti-TL2 Ab (10 µM), the combination of both, LPS (10 µM) alone, or LPS in combination with anti-TL2 Ab for 6 hours, as indicated. Mean ± SEM, *p < 0.05 vs. native, n = 3–4.

**Supplementary Figure 4.**
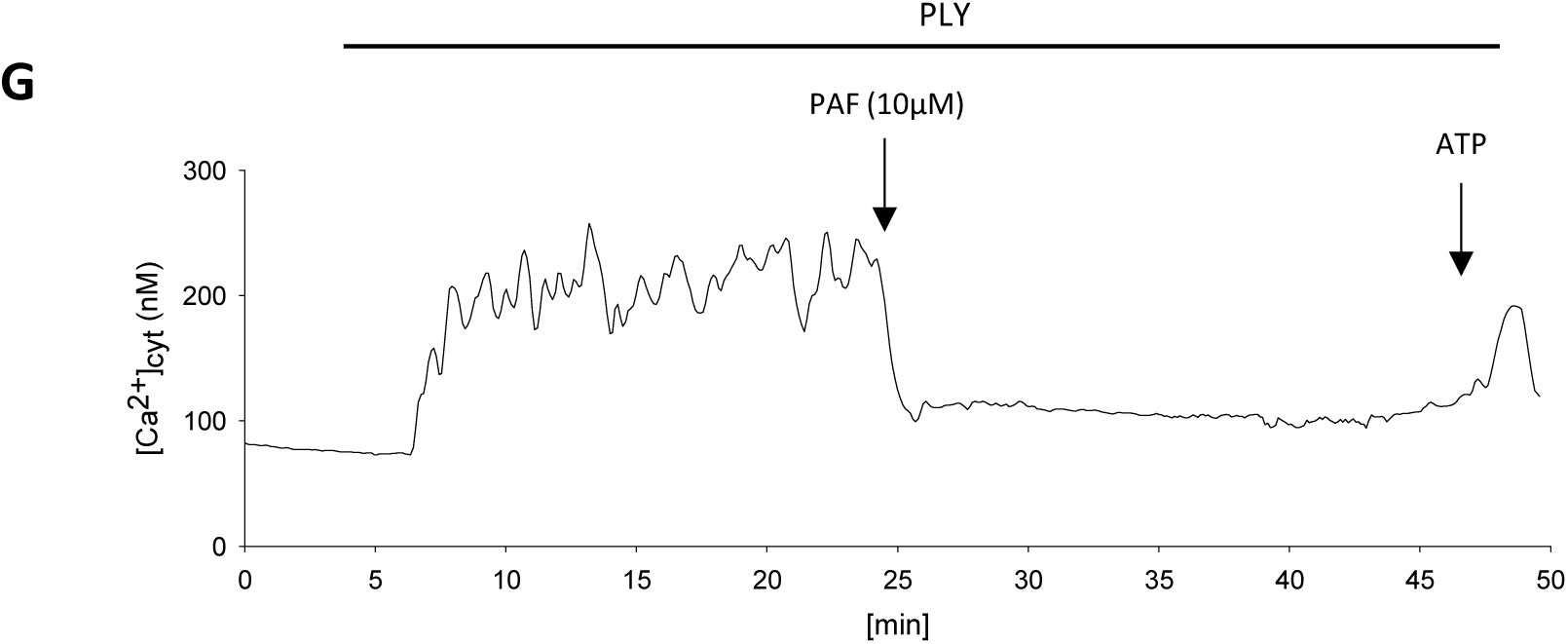
PAF inhibits PLY-induced Ca²⁺ signaling in HPMECs. **(A)** Tracings of endothelial [Ca²⁺]_cyt_ under baseline conditions and following sequential application of PLY (50µg/ml), PAF (10 µM), and ATP (100 µM) in the supernatant of untreated HPMECs, as indicated.

## Notes

### Competing Interest Statement

The authors have declared no competing interest.

